# Distinct cell states define the developmental trajectories of mucinous appendiceal neoplasms towards pseudomyxoma metastases

**DOI:** 10.1101/2022.05.26.493618

**Authors:** Carlos Ayala, Anuja Sathe, Xiangqi Bai, Susan M. Grimes, Jeanne Shen, George A. Poultsides, Byrne Lee, Hanlee P. Ji

## Abstract

Appendiceal mucinous neoplasms **(AMN)** are rare tumors of the gastrointestinal tract. They metastasize with widespread abdominal dissemination leading to pseudomyxoma peritonei **(PMP)**, a disease with poor prognosis. The cellular features of origin, differentiation and progression in AMN and PMP remain largely unknown. We characterized the distinct cell states of AMN and PMP using single-cell RNA-sequencing in 31 samples including AMNs, PMP metastases and matched normal tissues. We identified previously undescribed cellular features and heterogeneity in AMN and PMP. We discovered developmental cell states in tumor epithelial cells that ranged from progenitor phase to goblet cell differentiation, which was accompanied by elevated mucin gene expression. Metastatic PMP cells had a distinct cell state with increased mTOR and RAS signaling pathways and a series of upregulated cancer genes. We observed heterogeneity in a single PMP tumor as well as PMP metastases from the same patient. We validated our findings with immunohistochemistry, mass spectrometry on malignant ascites from PMP patients and gene expression data from an independent set of 63 PMPs.

## INTRODUCTION

Appendiceal mucinous neoplasms **(AMN)** are a rare indolent malignancy that originates in the appendix. AMNs can rupture resulting in intraabdominal dissemination. This condition is called pseudomyxoma peritonei **(PMP)** and is a cause of significant morbidity. Current treatment consists of cytoreductive surgery **(CRS)** and heated intraperitoneal chemotherapy **(HIPEC) (1)**. However, only 60% of PMP patients are eligible for this treatment **(2)**. The remainder are treated with systemic chemotherapy with limited benefit for most patients. The poor patient outcomes reflect the extent of metastasis and the specific cellular features of the tumor **(3)**.

There remain fundamental questions about the dysregulated biology of tumor cells in AMN and PMP. Determining these features may lead to improved therapeutic strategies.

AMN pathological specimens have been shown to contain goblet cell hyperplasia(4). Elevated mucin production, predominantly MUC2, is detected in AMN and PMP tissue. Some reports have stated that PMPs originate from MUC2-expressing goblet or goblet-like cells **(5)**.

However, PMP tumor epithelial cells do not have classic histopathologic characteristics of goblet cells **(6)**. Moreover, little is known about the pathways that distinguish AMNs from PMPs.

The definitive genomic studies of PMPs were conducted by Levine et al. (3, 7). They used conventional bulk gene expression analysis on the largest collection of appendiceal cancers to date. They identified genomic signatures that were distinct from colorectal cancer, confirming their unique biology. These gene expression signatures were indicative of oncogenic processes and pathways which were shown to have prognostic implications.

This study’s goal was to characterize the cellular and molecular features of AMN and PMP. PMPs have been challenging to study due to their low tumor cell cellularity and gelatinous nature. To characterize the gene expression of specific cell populations from both AMN and PMP, we applied single-cell RNA sequencing **(scRNA-seq)** in a discovery set, followed by independent validation approaches. Our analysis was focused on the origins and differentiation of the tumor epithelial cell populations. This study identified the cell states in tumor epithelium that were associated with the progression of AMN to PMP.

## RESULTS

### Study overview

This study included AMNs without perforation and PMPs of appendiceal origin. All specimens following surgery were reviewed by a board-certified pathologist at our institution to confirm the site of origin **(Table 1)**. The AMNs were low-grade appendiceal mucinous neoplasms **(LAMN)** restricted to the appendix without peritoneal dissemination. The PMPs were low-grade mucinous carcinoma peritonei **(LMCP)** or high-grade mucinous carcinoma peritonei **(HMCP)** of the appendix with peritoneal dissemination (**Supplemental Fig. 1A**). We used multiple approaches to characterize the cellular origins of this disease **(Fig. 1A)**. The first part involved scRNA-seq applied to a discovery cohort of 14 patients containing 31 samples from AMNs, PMPs and a subset of matched normal appendix or omental tissue. A subset of the PMP samples represented multiple metastatic sites from the same patient. Tumor sites included the omentum, small bowel mesentery, liver capsule, and ovaries.

**Table 1.** AMN and PMP samples.

| **Tumor type** | **Patient ID** | **Tissue sites for single-cell analysis** | **No. of samples for single-cell analysis** | **Mass spectrometry on ascitic fluid** | **Diagnostic cancer gene sequencing** |
| --- | --- | --- | --- | --- | --- |
| AMN | P8139 | Appendix tumor | 1 | No | NA |
| AMN | P8142 | Normal appendix | 2 | No | NA |
| AMN | P8153 | Appendix tumor and normal | 2 | No | NA |
| AMN | P8333 | Appendix tumor and normal | 2 | No | NA |
| AMN | P8617 | Appendix tumor and normal | 2 | No | NA |
| PMP | P8336 | Peritoneum tumor | 1 | No | NA |
| PMP | P5853 | Peritoneum tumor | 1 | Yes | *KRAS* G12D |
| PMP | P8587 | Omentum tumor | 1 | No | NA |
| PMP | P8605 | Appendix mesentery, omentum and ovary tumors | 3 | Yes | *GNAS*, *KRAS* |
| PMP | P8606 | Appendix and small bowel mesentery tumors, normal omentum | 3 | No | NA |
| PMP | P8620 | Appendix and omentum tumors, normal omentum | 3 | No | NA |
| PMP | P8629 | Appendix mesentery and small bowel mesentery tumors | 2 | Yes | *KRAS* G12D, *GNAS* |
| PMP | P8639 | Appendix, omentum, ovary and appendix mesentery tumors | 4 | No | NA |
| PMP | P8642 | Appendix mesentery, omentum, liver capsule tumors, normal omentum | 4 | No | NA |

AMN: Appendiceal mucinous neoplasm, PMP: pseudomyxoma peritonei, NA: Not Available

**Figure 1.**
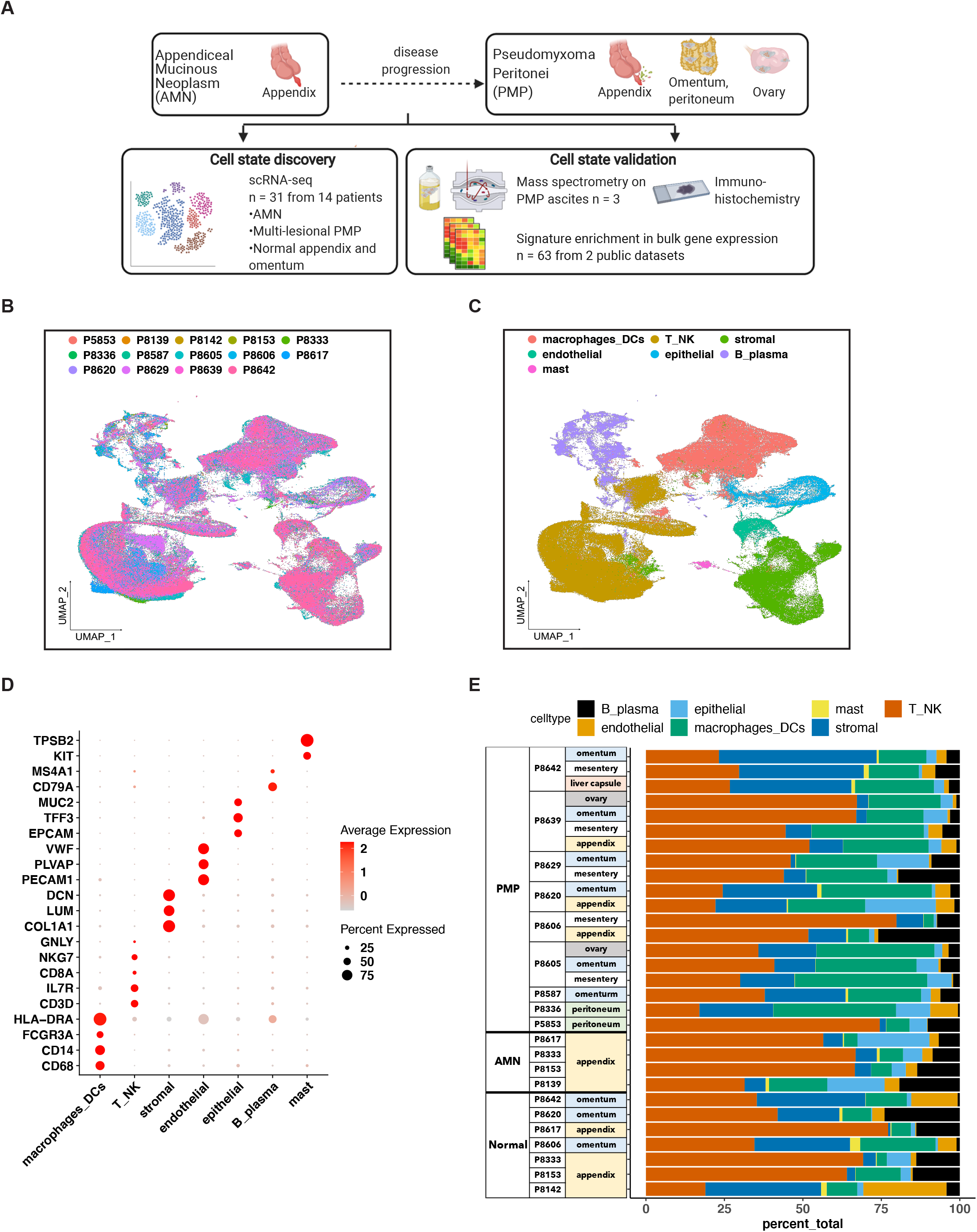
(A) Schematic representation of study design. (B-C) UMAP representation of dimensionally reduced data following batch correction and graph-based clustering of all datasets annotated by (B) samples and (C) cell lineages. (D) Dot plot depicting average expression levels of specific lineage-based marker genes together with the percentage of cells expressing the marker. (E) Proportion of cell lineages detected from each sample annotated by normal, AMN and PMP designation.

We validated our findings for specific cell populations using immunohistochemistry **(IHC)**. To determine if critical gene expression markers could be detected as secreted proteins, we conducted mass spectrometry on ascites samples from three PMP patients. Using an independent cohort of 63 PMP tumors, we confirmed the enrichment of cellular states identified in our discovery cohort.

### Single cell RNA-seq profiling of AMN and PMP tumors

Samples included normal appendix, normal omentum, AMNs or PMPs. The scRNA-seq analysis from all samples provided a total of 299,718 cells **(Supplemental Table 1)**. We applied the Harmony algorithm to correct for batch variance in the data (8). We observed that each cluster had cells from different patient tumors, indicating adequate removal of noise from experimental batch effects **(Fig. 1B)**. We verified this computationally by calculating a similarity metric called the Adjusted Rand Index **(ARI)** (9). Comparison of cluster assignments with experimental batch had an ARI of 0.05. The low, near-random value confirmed that cluster assignments were the result of intrinsic tumor cell properties and not unduly influenced by batch effects.

For each cluster, we assigned cell lineages based on the expression of established marker genes **(Fig. 1C - D)**. This identified epithelial (*EPCAM, TFF3, MUC2*), stromal fibroblasts (*DCN, COL1A1, LUM*), endothelial (*VWF, PLVAP, PECAM1*), T (*CD3D, IL7R, CD8A*), NK (*NKG7, GNLY*), B or plasma (*MS4A1*, CD79A), mast (*TPSAB1, KIT*) and macrophage or dendritic cells (*CD68, CD14, FCGR3A, HLA-DRA*). All lineages were identified in varying proportions across all samples **(Fig. 1E)**. Next, we performed a secondary clustering analysis with batch-correction for each cell lineage, enabling the detailed characterization of its transcriptional phenotypes.

### Delineating the epithelial cell subtypes in AMNs and PMPs

We focused on determining the characteristics of the different epithelial subtypes present in these tumors. We conducted an independent clustering analysis of the epithelial cells from the AMNs, PMPs and normal appendix containing at least 10 cells **(Supplemental Fig 1B, Supplemental Table 2)**. This threshold was required for evaluating cell states and differential expression. We compared the gene expression of each individual cell to a reference single cell atlas of mucosal cell lineages of the human large intestine (10). This step relied on the SingleR method which can identify cell types in a query dataset using cell-type specific reference transcriptomic datasets **(Supplemental Methods)** (11).

Next, we verified cell assignments by examining the expression of specific lineage marker genes. We identified enterocytes (*CA2, SLC26A2, FABP1*), stem (*LGR5, ASCL2, OLFM4*) and goblet cells (*MUC2, TFF3, SPINK4, CLCA1, SPDEF, FCGBP*) **(Fig. 2A)** throughout all samples. Rare cell types included chemosensory tuft (*POU2F3, LRMP, TRPM5*) and neuroendocrine cells (*CHGA*).

**Figure 2.**
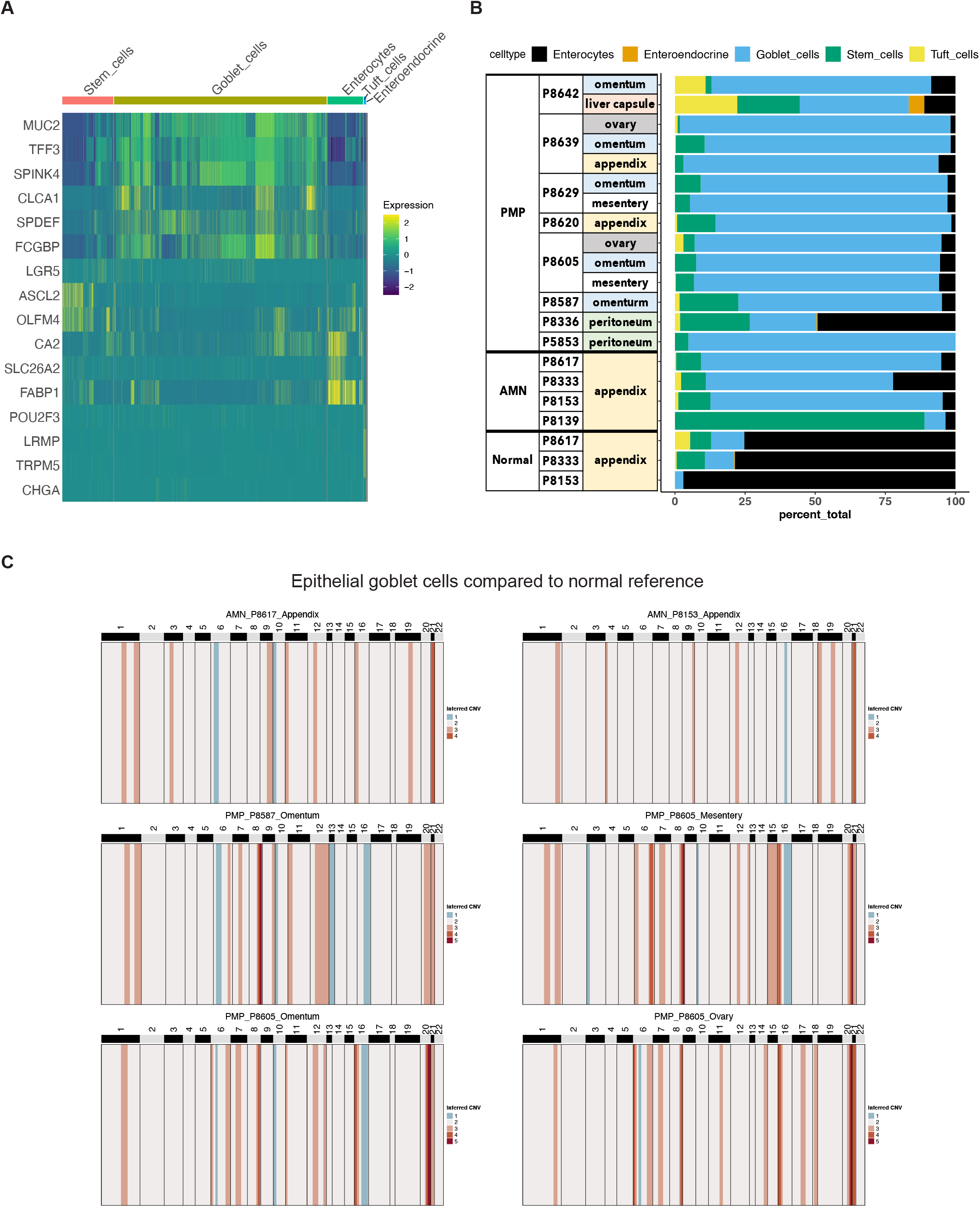
(A) Heatmap depicting expression of lineage marker genes following reference-based assignment of tumor and normal epithelial cells. (B) Proportion of epithelial cell lineages detected from each sample annotated by normal, AMN and PMP designation. (C) Heatmap representation of inferred single-cell CNV profiles of goblet tumor epithelial cells per respective sample compared to reference cells from normal epithelium, immune and stromal cells. CNV legend: state 1 = complete loss, state 2 = loss of one copy, state 3 = neutral, state 4 = addition of one copy, state 5 = addition of two copies, state 6 = addition of more than 2 copies.

Using this reference transcriptome-based mapping approach, we identified goblet cells as the major epithelial subtype in both AMN and PMP tumors **(Fig. 2B)**. To compare the composition of normal tissue versus AMN or PMP, we conducted a statistical analysis for differential cell type abundance using a generalized linear model **(Supplemental Methods)** (12, 13). Compared to normal appendiceal tissue, goblet cells were significantly more abundant in PMPs (adjusted P value 2.49E-50; Wald test) and AMNs (adjusted P value 1.21E-33). Meanwhile, stem cells were significantly enriched in AMN compared to normal tissue (adjusted P value 2.65E-17) or PMP (adjusted P value 1.93E-100). Notably, this might have been influenced by P8139 for which the majority of the cells had a molecular profile of intestinal stem cells. We analyzed the expression of *MUC2*, encoding for the predominant extracellular mucin protein detected in AMN and PMP.

This protein is expressed in cells of the gastrointestinal tract, but not the omentum (4). Compared to normal appendix cells, the AMN and PMP epithelial cells showed significantly higher *MUC2* expression, with the highest expression in PMP cells (ANOVA FDR P-value < 2.2E-10) **(Supplemental Fig. 1C)**. We observed a similar overexpression pattern in PMP cells for a gene signature encoding gel-forming mucins (*MUC2, MUC5B, MUC5A, MUC6, MUC19*) (14) **(Supplemental Fig. 1D)**. Altogether, our results identified the presence of cells with a goblet cell signature in patients with peritoneal carcinomatosis from appendiceal origin.

### Genomic instability in neoplastic goblet cells from AMNs and PMPs

To confirm that these epithelial cells were neoplastic, we characterized the extent of somatic copy number variations **(CNVs)** using the inferCNV program (15, 16) **(Supplemental Methods)**. Briefly, CNVs were estimated based on average gene expression of a 10Mb or larger windowed segment. These delineated allelic imbalances covering chromosome arms. The CNVs present in goblet cells from AMN and PMP tumor epithelium were compared to normal appendiceal epithelial cells and other randomly sampled non-epithelial cells in the TME. These normal cells lacked CNVs.

In contrast, the goblet epithelial tumor cells from all AMNs and PMPs had multiple CNVs **(Fig. 2C, Supplemental Fig. 2)**. Large CNVs with imbalances of entire chromosome arms are indicators of chromosomal instability. Altogether, their presence identified these goblet epithelial cells as neoplastic.

A gain of chromosome 9 and 21 was a frequently observed CNV across all samples including both AMNs and PMPs. Interestingly, aberrant activation of the NOTCH 1 receptor located on chromosome 9 has been described in PMP (17). We also observed gains in chromosomes 1q and 20q that have been previously identified in PMP (17, 18).

Among PMP tumors in different anatomic locations from the same patient, we identified metastatic clonal divergence. For example, P8639 had three metastases, all with a gain of chromosomes 1, 9 and 21 **(Supplemental Fig. 2)**. However, only the appendiceal metastasis had a gain of chromosome 15. P8629 had two metastases: only the mesentery lesion had a gain of chromosomes 12 and 15 compared to the omentum site **(Supplemental Fig. 2)**. P8605 had three metastatic sites **(Fig. 2C)**. All had gains in chromosomes 1, 8, 20 and 21. Only the mesenteric site had a gain in 15. This result confirmed that clonal heterogeneity was present among the different tumor sites from the same patient.

### Developmental trajectory and cell states among AMN and PMP epithelium

Cell differentiation is a hallmark of cancer progression such as what may occur in AMNs as they progress to PMPs. Using single-cell gene expression data, one can identify a ‘trajectory’ or path for cells along this differentiation process. Cells are positioned along this trajectory as a measure of ‘pseudotime’. We analyzed the pseudotime trajectories and defined cell states from epithelial cells derived from normal appendix, AMN and PMP using the Monocle algorithm (19). Subsequently, we evaluated the specific features of each state.

The normal appendiceal epithelium defined the root of the trajectory, which we named State 0 **(Fig. 3A)**. Branching from the root, we identified five additional cellular states **(Fig. 3A, B)**. States 1 and 2 had lower pseudotime values; indicative of early differentiation and representing progenitor cells. States 3, 4 and 5 had higher pseudotime values, indicative of later stages of development.

**Figure 3.**
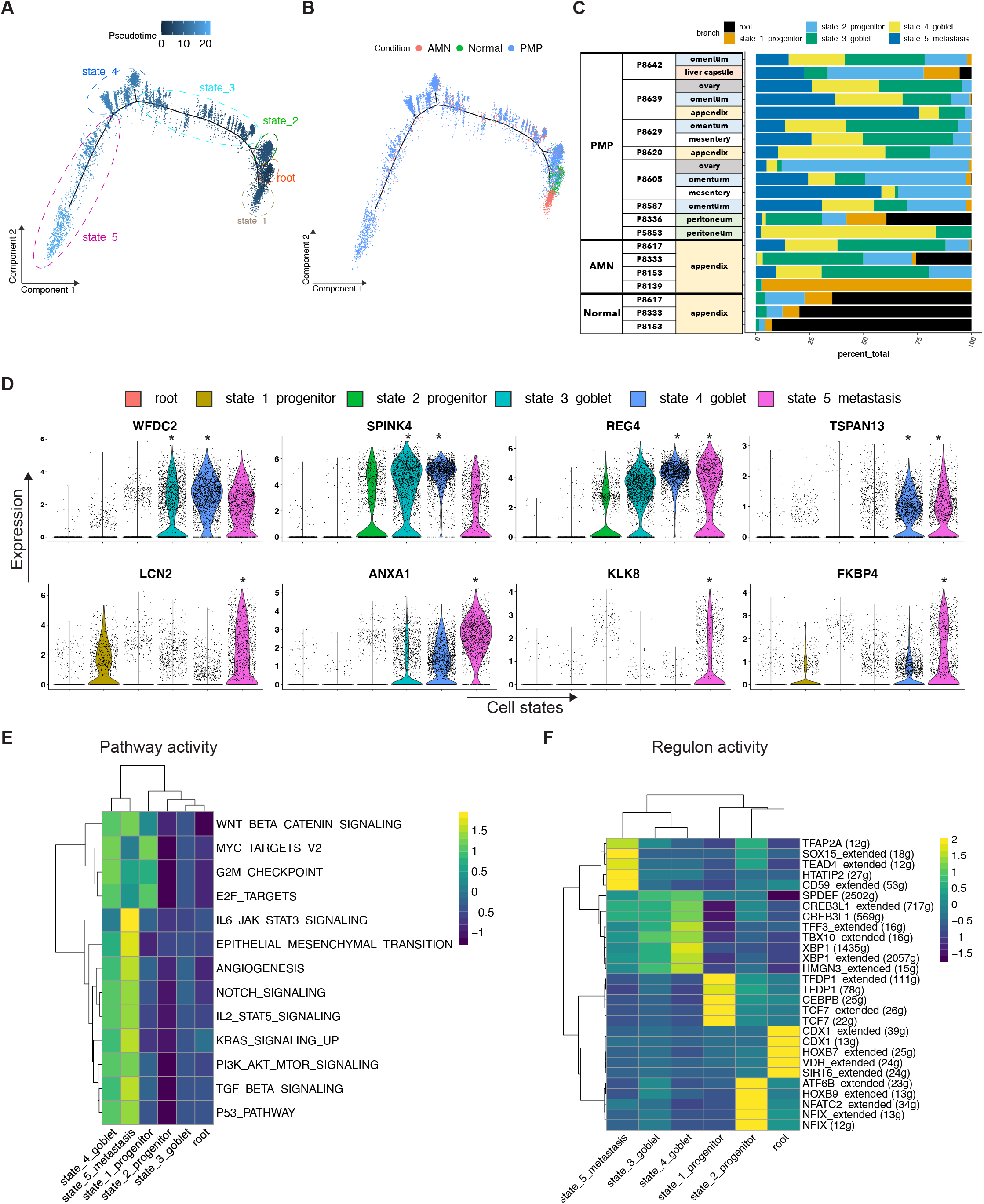
(A-B) Trajectory plots of epithelial cells colored by their (A) pseudotime values and additionally annotated by state, (B) condition. (C) Proportion of trajectory states within epithelial cells detected from each sample annotated by normal, AMN and PMP designation. (D) Violin plots indicating expression of selected genes in each trajectory branch. * indicates cell state with significant expression (adjusted p-value < 0.05) for that gene. (E) Heatmap depicting average GSVA enrichment score of selected Hallmark pathways per trajectory branch (ANOVA Tukey HSD p-value < 0.05). (F) Heatmap depicting average Area Under the Curve value of respective regulon per trajectory branch (ANOVA Tukey HSD p-value < 0.05). Numbers in parentheses indicate the number of genes in the examined gene regulatory network.

We measured the fraction of epithelial cells for each cellular state across samples **(Fig. 3C, Supplemental Table 3)**. State 1 was over-represented by P8139 likely representing a unique biological trait of this patient compared to the remainder of the AMN and PMP cohort. Using differential abundance analysis **(Supplemental Methods)**, we determined that State 3 was most common among AMN compared to normal and PMP (adjusted P value <= 1.71E-7; Wald test). States 4 and 5 were significantly more common among PMP versus AMN (adjusted P value <= 4.62E-15). For an individual tumor, multiple cell states were present, indicative of intra-tumor heterogeneity. For example, all the PMP tumors had subpopulations from all four differentiation states in varying proportions. In summary, our results identify the presence of multiple cell states in lesions of AMN and PMP patients. Additionally, states 2 and 3 were dominant in AMN while State 5 was prevalent in PMP. Altogether, the trajectory analysis results support the notion of a cellular differentiation spectrum from AMN to PMP pathology.

### Genes defining each cell state

To elucidate cell state properties in greater detail, we performed differential expression and pathway analysis. The root or State 0 expressed enterocyte markers (*FABP1, GUCA2A, CA2, ANPEP*, etc.) **(Supplemental Table 4)**, indicative of normal epithelial cells found in the appendix. State 1 had high expression of the stem cell marker *OLFM4* as well as chemokines *CCL20, CXCL3, CXCL14* and *CXCL1*. State 1 was associated with P8139’s AMN tumor and indicated an inflammatory progenitor cell state, which likely represents a unique biological trait of unknown clinical significance.

State 2 cells were present in the remaining AMN tumors and majority of PMP tumors. This state had a gene expression signature associated with progenitor cells **(Supplemental Table 4)**.

This progenitor cell state included high expression of *SOX4* that regulates stem cell differentiation (20) and *ELF3* that promotes tumor growth (21). These transcription factors have been associated with intestinal and goblet cell lineage differentiation, respectively (22, 23). This state also demonstrated high expression of *SPDEF* and *MUC5AC* which regulate goblet cell phenotype (24). *SPDEF* is an essential transcriptional factor that is involved in goblet cell differentiation (24). Overall, these results suggest that the acquisition of goblet cell features is an early developmental event.

State 3 was enriched in AMN cells and State 4 was enriched in PMP cells **(Fig. 3C)**. Compared to the other cell states, they significantly overexpressed genes indicative of terminal goblet cell differentiation including*TFF3, MUC5AC, MUC2, SPINK4, WFDC2* and *FCGBP* **(Fig. 3D, Supplemental Table 4)**. Several genes were specific to State 4 including *REG4* and tetraspanin genes such as *TM4SF1, CD9, CD63, TSPAN8, TSPAN13*. These genes are contributors to proliferation and metastasis in other malignancies (25, 26).

State 5 was widely represented among the PMPs compared to the AMNs. The differentially expressed genes included *ANXA1, FKBP4, LCN2, VNN1, TCN1, TACSTD2, DUOXA2, CAECAM5, CAECAM6* and members of the Kallikrein **(KLK)** family (*KLK6, KLK8, KLK10* and *KLK11*) **(Fig. 3D, Supplemental Table 4)**. State 5’s gene expression signature was present in 11 of the PMP tumors **(Supplemental Fig. 3A, B)**. These genes have been shown to regulate cancer metastasis and are associated with poor prognosis (27-32). However, they have not been described as playing a role in PMPs in prior studies using bulk RNA analysis.

For example, *ANXA1* is a phospholipid-binding protein that regulates cancer cell proliferation and metastasis (33). *KLK8* is highly expressed in several tumor types including pancreatic and colorectal cancer. In mouse models, *KLK8* overexpression had pro-proliferative and anti-apoptotic effects (34). *FKBP4* belongs to the immunophilin family with roles in immunoregulation, protein folding and trafficking. It is overexpressed in many tumor types with an uncharacterized role in cancer (35). These genes are candidate markers distinguishing the metastatic state of PMPs and may play a role in their cancer biology.

### Biological pathways and regulatory genes defining epithelial cell states

For each cell state, we conducted a biological pathway analysis with the Gene Set Variation Analysis **(GSVA)** tool. The pathway annotation was based on the Hallmark gene sets – these groups of genes define specific biological pathways and can be assigned to a given cell state (36, 37).States 1 (progenitor phenotype), 4 (goblet cell differentiation) and 5 (PMP metastasis) had significant upregulation of WNT pathway involved in stem cell maintenance, MYC signaling and cell cycle pathways (ANOVA with Tukey HSD p-value <= 3.6e-04) **(Fig. 3E)**.

States 4 and 5 were also notable for the upregulation of cancer biology pathways including: PI3K-mTOR, p53, KRAS and Notch signaling (ANOVA with Tukey HSD p-value <= 7.46e-06) **(Fig. 3E)**. State 5 had significant upregulation of pathways associated with cancer development and progression that included epithelial mesenchymal transition **(EMT)**, TGF-β, JAK-STAT and angiogenesis signaling (ANOVA with Tukey HSD p-value <= 4.2e-08). These pathways thus support the development of PMP metastasis. Importantly, *KRAS* represents a common mutations found in AMN and PMP (18). Meanwhile, NOTCH pathway activation has been shown to promote goblet cell differentiation(38). Dysregulation in TGFB, WNT, MY and EMT pathways overlap with what Levine et al. described in their gene expression studies (3, 7).

Next, we identified the regulatory genes that defined the transcriptional characteristics of these cell states using the SCENIC algorithm (39) **(Fig. 3F)**. These regulatory genes or regulons represent transcription factors that regulate the expression of downstream target genes.

Progenitor states 1 and 2 had significantly increased activity of *TCF7* and *NFIX* regulons (ANOVA with a Tukey HSD p-value <= 2.68e-11). *TCF7* was recently discovered as a regulator of goblet cell fate using gut derived organoids(40). High expression of *NFIX* has previously been noted in gut stem cells (41). States 3 and 4 indicative of goblet cell differentiation had significantly high activity of *SPDEF*. This transcription factor regulates differentiation into goblet cells (24). Meanwhile, *SPDEF* activity was lower in State 5, relative to State 3 and 4. This state 5 was associated with EMT related expression. This result suggests an early acquisition of *SPDEF* activity that promotes goblet cell differentiation, followed by additional regulatory changes associated with progression to PMP. We also noted increased regulatory activity of *TFAP2A, TEAD4* and *CD59* **(Fig. 3F)**. These genes have been shown to promote EMT in lung and colon cancer, among other malignancies (42-44). Overall, the pathway and regulon analysis identified a series of regulatory steps that are associated with stem cell activity, goblet cell differentiation and metastatic progression in AMN and PMP.

### Heterogeneity among multiple site PMP metastases and their cell states

Using the results of the trajectory analysis, we examined the cellular heterogeneity for a subset of patients with multiple metastatic samples. We used three PMP patients (P8605, P8629, P8639) that met criteria with an adequate number of cells (10 or greater) for this analysis. We previously observed clonal divergence among these samples based on CNV alterations (**Fig. 2 and Supplemental Fig. 2**). Therefore, it was of interest to see if the cell states and gene expression also had displayed heterogeneity.

Each metastasis from the same patient had varying proportions of the different cell states **(Fig. 3C)**. For example, P8605’s ovarian metastasis had a higher proportion of cells in State 2 compared to the mesenteric site. The inverse was observed for State 5 with higher proportion in mesenteric versus the ovarian metastasis.

We examined the gene expression differences among the metastatic sites. Each site from P8605 had distinct gene expression signatures. For the mesenteric metastasis, we observed significantly increased expression of metastasis-related genes *ANXA1* and *KLK8* compared to the omental and ovarian implants (adjusted p-value < 0.05) **(Supplemental Fig. 4A)**. The omental metastasis had higher expression of the goblet cell differentiation markers *FCGBP* and *TFF3*. The ovarian site had increased expression of the progenitor state marker *SOX4*. The three lesions also differed in their expression of *GNAS*, a frequently mutated gene in PMP that has been demonstrated to regulate mucin production (45). This results indicates the divergence in clonal properties across metastatic sites.

For patient P8629, the omental metastasis had a higher expression (adjusted p-value < 0.05) of goblet cell differentiation markers *TFF3, FCGBP, MUC2* and *SPINK4* **(Supplemental Fig. 4B)**. Conversely, the mesenteric lesion had enriched expression of the progenitor marker *ELF3* and *GNAS*. The tumors from P8629 and P8605 underwent diagnostic cancer gene sequencing.

Both patient tumors had mutations in *GNAS* (**Table 1**). Importantly, our analysis discovered a variation in *GNAS* expression level across multi-lesion metastasis in the same patient.

Patient P8639 had two metastatic sites from the omentum and ovary together with the primary appendiceal cancer. The metastatic sites had differential expression of the following genes: *TCN1* associated with poor prognosis and metastasis in colon cancer (28); goblet cell marker *WFDC2*; and *GNAS* **(Supplemental Fig. 4C)**. Altogether, these results confirm that PMP metastatic sites from the same patient have different properties of stemness, goblet cell identity and metastasis differentiation.

### Protein markers present in PMP tumors

One of our major findings was that PMP epithelial cells have cell states defined by goblet cell and metastasis differentiation. To validate specific markers of these cell states, we used a combination of immunohistochemistry **(IHC)** and mass spectrometry on a subset of the PMP tumors and ascites respectively. The goal with both approaches was the detection and identification of proteins that could be correlated with our transcriptomic data for the neoplastic cells. For the IHC study, we used antibody markers that defined goblet cell state (MUC2 and AGR2) and PMP metastasis State 5 (ANXA1). Mass spectrometry enabled an unbiased assessment of proteins detected in ascites from PMP patients.

For the three PMP tumors (P8629, P8639 and P8605), MUC2 and AGR2 demonstrated intense staining among the epithelial tumor cells **(Fig. 4A)**. Likewise, ANXA1 was detected in all PMPs albeit with less intense staining than MUC2. The tissues tested included mesentery and ovary, which do not contain goblet cells stained by MUC2 and AGR2. Their detection therefore represents the identification of neoplastic cells of gastrointestinal origin that have metastasized. Altogether, our results show that all markers were expressed among the PMP tumor epithelial cells.

**Figure 4.**
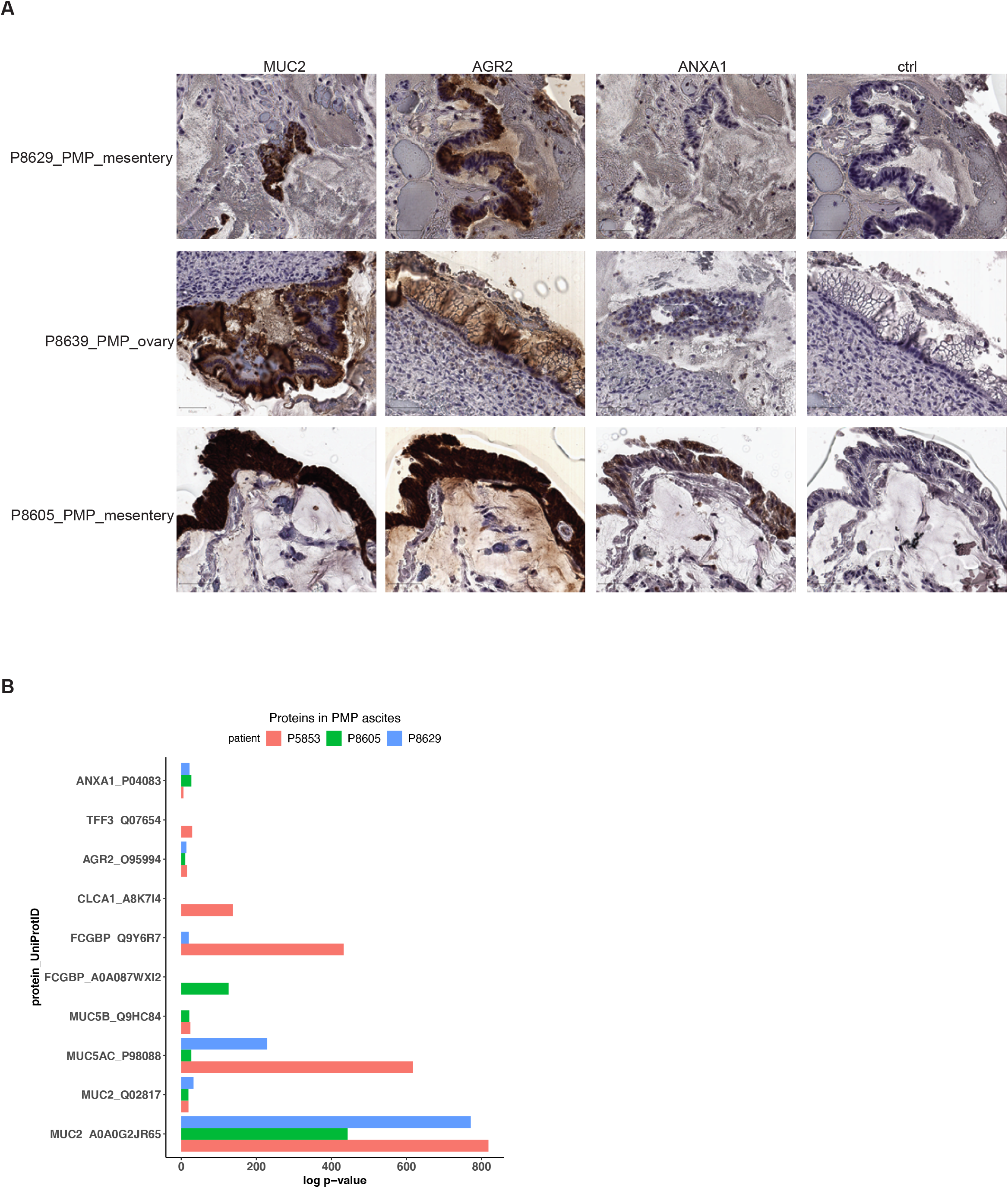
(A) Representative images of immunohistochemical staining of respective tumor samples for MUC2, AGR2, ANXA1 or representative isotype control. Scale bar = 50 µm. (B) Selected proteins detected in mass spectrometry of PMP ascites samples with their detection p-value.

Malignant ascites samples were available from three PMP patients (P5853, P8605 and P8629). Goblet cells are secretory cells of the internal lumen of the gastrointestinal tract. We determined if products of goblet cells, normally found inside the gastrointestinal lumen following secretion, were secreted into the ascites using mass-spectrometry. Using a probabilistic model, we identified the likelihood that peptide spectrum matches arise by random chance (46). Next, we used a confidence threshold requiring a detected protein to have a log p-value at least 2.0 fold lower than the log p-value of the top decoy protein (47). Overall, we detected 299 proteins in P5853, 300 in P8605 and 276 in P8629 **(Supplemental Table 5)**.

Across all samples, we observed the expression of mucin proteins MUC2, MUC5AC and MUC5B, goblet cell markers AGR2 and FCGBP and metastasis State 5 marker ANXA1 **(Fig. 4B, Supplemental Table 5)**. P5853’s ascites had goblet cell secretory products TFF3 and CLCA1 (48). In summary, this protein analysis supported a goblet cell identity for the PMP tumors. We detected goblet cell secretory products in ascites that are encoded by the transcriptome of neoplastic cells in non-gastrointestinal tissue (e.g. omentum, mesentery, ovary). More importantly, our results demonstrated a direct role for these cells in the secretion of their intracellular products into malignant ascites.

### Evidence of PMP cell states in independent cohorts

We validated our scRNA-seq findings using independent gene expression data (Affymetrix microarrays) from 63 PMPs (3). We performed gene set variation analysis **(GSVA)** (36) on this expression data. For this analysis, we used the expression signatures representing significantly differentially expressed genes per cell state **(Fig. 5A)**. We aggregated states that were closely related. States 1 and 2 represented the progenitor state, 3 and 4 represented the goblet cell state. We examined PMP metastasis State 5 and root State 0 separately. The GSVA analysis provided a quantitative enrichment score, which indicated whether the PMPs expressed the gene signature of any given cell state.

**Figure 5.**
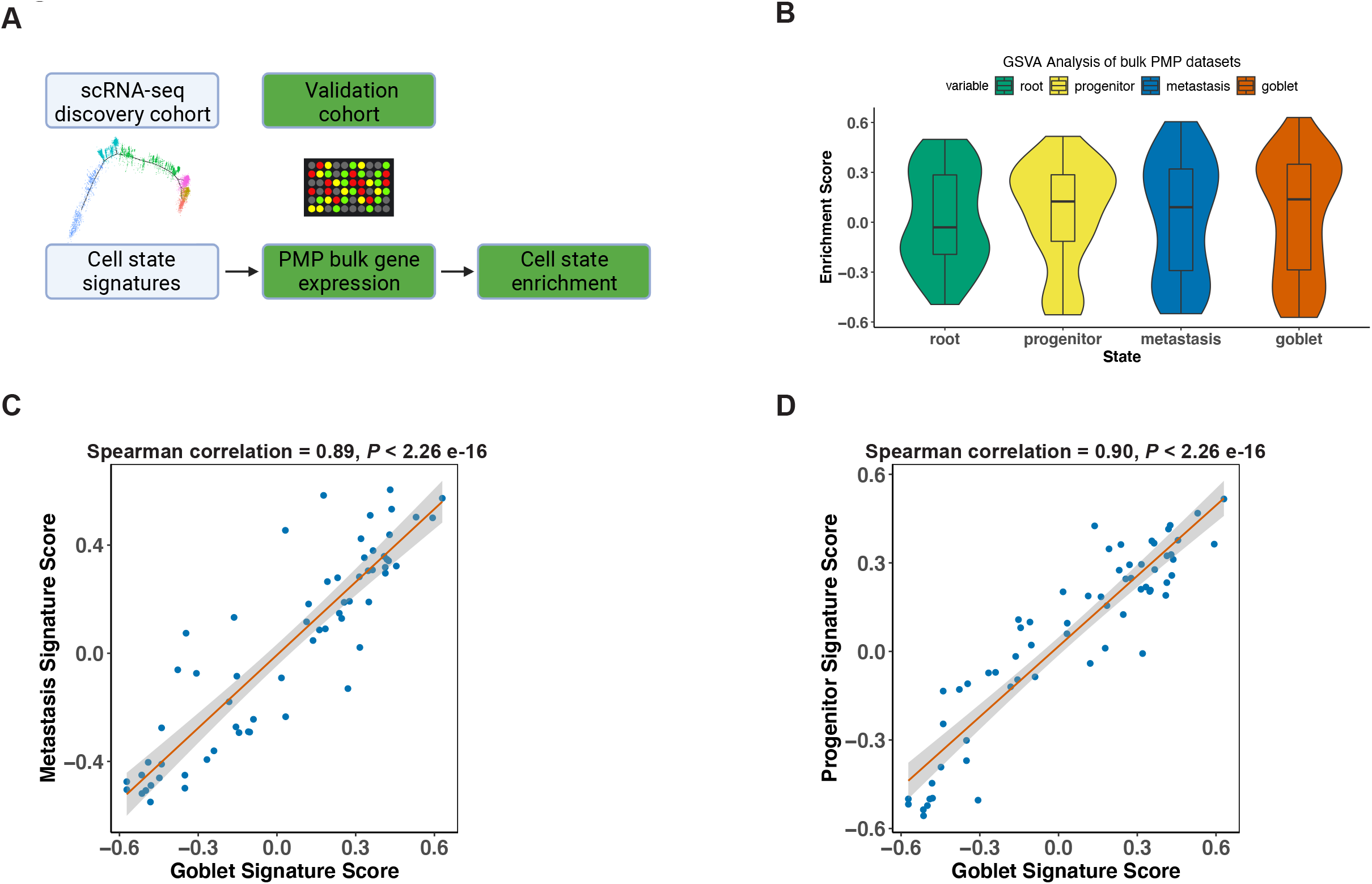
(A) Schematic representation of signature enrichment analysis. (B) GSVA scores for respective tumor epithelial cell state in validation bulk dataset samples. (C-D) Scatter plots showing Spearman correlation of respective tumor epithelial cell state signatures across all samples from the validation bulk datasets.

The majority of samples had positive GSVA enrichment scores for the root state (49.2%), progenitor state (61.9%), goblet cell differentiation state (58.7%) and the metastatic state (57.1%) **(Fig. 5B)**. Hence, cell states from our single-cell discovery cohort were detected in independent PMP cohorts. The low cellularity of these tumors makes it challenging to detect tumor epithelium specific features from conventional bulk sequencing assays. This result indicated that having identified granular cell states using scRNA-seq, we can identify them in bulk datasets using their signatures. Furthermore, the goblet state enrichment scores were significantly correlated with metastasis **(Fig. 5C)** and progenitor **(Fig. 5D)** scores across all samples. This result confirmed our finding from the discovery cohort that heterogenous cell states co-exist within a PMP tumor.

### Infiltrating lymphocytes in the AMN and PMP microenvironment

We characterized the lymphocytes in the TME of the AMNs and PMPs. Our major focus was determining if these tumors contain infiltrating lymphocytes. Initially, we annotated lymphocytes by comparing individual cell transcriptomes to a reference pan-cancer atlas of tumor infiltrating immune cells and verified assignments based on the expression of marker genes **(Supplemental Methods)** (49).

We identified B and plasma cells (*MS4A1, CD19, etc*.) **(Fig. 6A)**. We detected T cell subsets including naïve-like (*CCR7, SELL*), regulatory T **(Tregs)** (*FOXP3, IL2RA, etc*.) and follicular helper-like cells **(TFh-like)** (*CXCL13, TNFRSF18, CTLA4, etc*.). We identified cytotoxic CD8 T (*CD8A, CD8B, CCL5, etc*.), activated-exhausted CD8 T cells (*CD8A, CD8B, CXCL13, PDCD1, CTLA4, etc*.) and NK cells (*KLRD1, KLRF1, GNLY, etc*.) (49, 50). Cell subsets were detected across normal, AMN and PMP samples (**Fig. 6B**). This indicates that AMN and PMP tumors contain a lymphocytic infiltrate. A recent study in PMP using spatial gene expression analysis supports this finding (51). While the absolute number of cells varied by sample **(Supplemental Table 6)**, all tumors contained TFh-like CD4 T, cytotoxic CD8 T and NK cells. Very few cells mapped to the exhausted CD8 phenotype.

**Figure 6.**
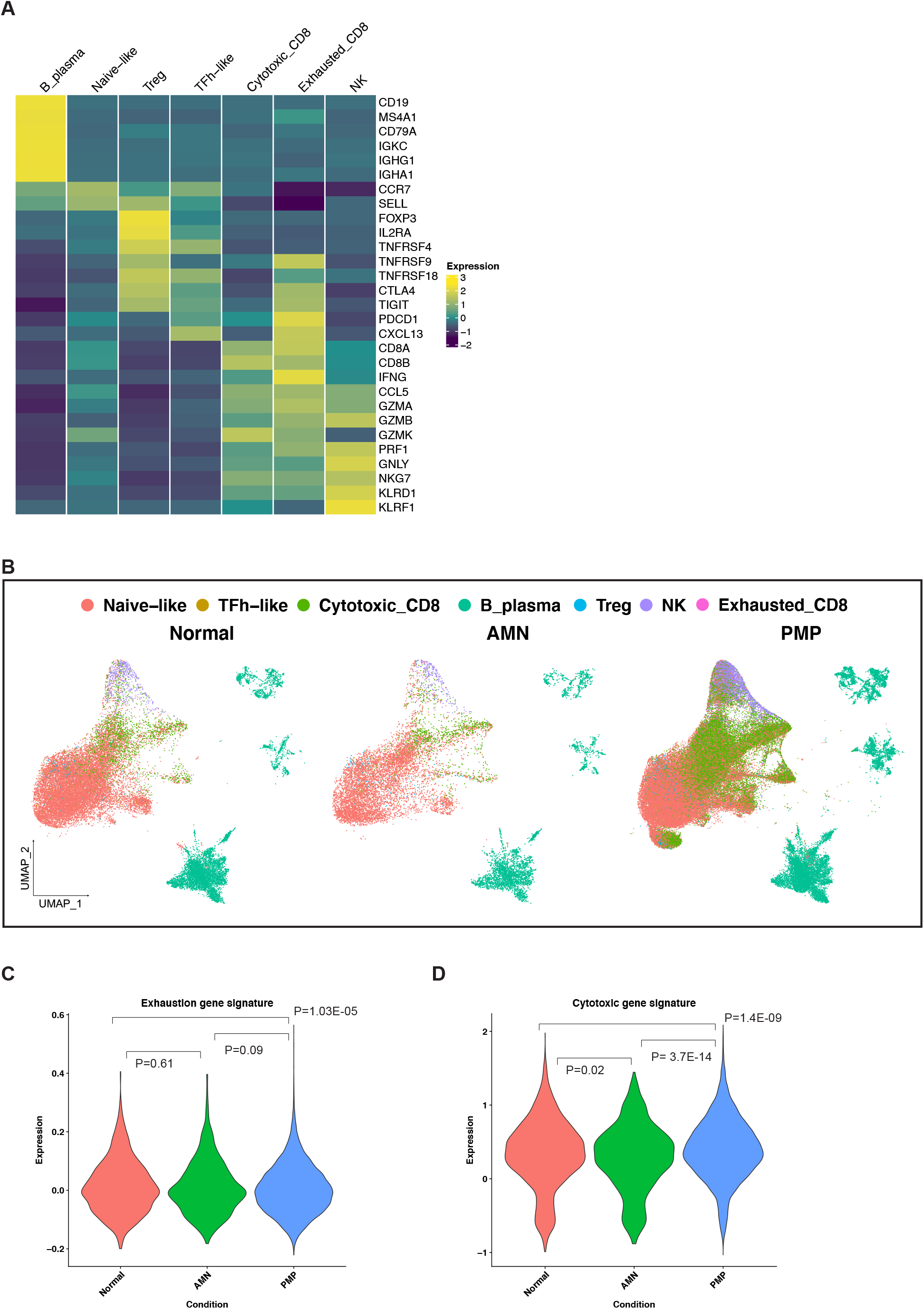
(A) Heatmap depicting average expression levels of respective genes in lymphocyte subsets from normal, AMN and PMP samples. (B) UMAP representation of dimensionally reduced data following batch correction and graph-based clustering of all datasets annotated by lymphocyte subsets across normal, AMN and PMP samples. (C-D) Violin plots depicting expression of (C) exhaustion or (D) cytotoxicity gene signature with ANOVA Tukey HSD p-value in normal, AMN and PMP infiltrating CD8 T cells.

The appendix and omentum are lymphocyte rich organs. Infiltrating CD8 T cells in AMN and PMP could indicate either anti-tumor cells or bystander cells resembling those in normal tissue. Infiltrating lymphocytes in AMN and PMP largely co-clustered with those in normal tissue (**Fig. 6B**), indicative of transcriptional similarity. An increase in exhaustion or dysfunction has been linked to CD8 T cells that recognize tumor antigens (50, 52). We evaluated the expression of an exhaustion associated gene signature in these cells (53) (**Supplemental methods, Supplemental Table 7**). CD8 T cells in AMN had no significant changes in exhaustion gene signature expression compared to CD8 T cells in normal tissues, while those in PMP had a reduced expression (**Fig. 6C**). This indicated that the CD8 T cell infiltrate in tumors is not specific to TME induced reprogramming. This finding further supported similarity of tumor infiltrating CD8 T cells with bystander cells in normal tissue. We next assessed the expression of a gene signature of cytotoxic effector genes in these CD8 T cells (16, 54) (**Supplemental methods, Supplemental Table 7**). CD8 T cells in AMN had a reduction in cytotoxic gene expression compared to normal tissue (**Fig. 6D**). Cells infiltrating PMP had increased cytotoxic gene expression. Hence, these infiltrating CD8 T cells in PMP continued to have high cytotoxic potential. Overall, these results suggest that the AMN and PMP TME is not “cold” and contains infiltrating lymphocytes. While these cells lacked features suggestive of anti-tumor activity, their cytotoxic potential could be harnessed further for strategies that aim to convert bystander cells into infiltrating cells that can contribute to anti-cancer activity (52). This is further supported by recent studies that demonstrated an *ex vivo* T cell activation in PMP tumors (55-57).

## DISCUSSION

The cellular characteristics of AMNs and their progression to PMP have been a challenge to study due to their rarity, low cellularity and complex gelatinous composition. The epithelial cell population is often diluted among the other cell types and mucinous components. To overcome these challenges, we used scRNA-seq on a set of appendiceal mucinous neoplasms without peritoneal dissemination (AMNs) and with peritoneal dissemination (PMPs). Given the low cellularity of these tumors, we combined low- and high-grade PMP lesions in our analysis. Our results identified the cellular features of low abundance tumor epithelial cells present in gelatinous appendiceal tumors with and without peritoneal dissemination.

An important aspect of our study was determining the origin and trajectory of AMN/PMP neoplastic cells. The discovery of MUC2 expression implicated an enteric rather than ovarian origin for PMP **(4)**. However, prior studies did not firmly establish whether this was due to acquisition of MUC2 expression by non-gastrointestinal tumor cells and/or cells with an intestinal lineage origin. Our analysis conclusively established a goblet cell identity of intestinal origin for the first time. Moreover, we observed tumor cells with goblet cell features in peritoneal metastases. These tissues normally lack intestinal epithelial cells (e.g. omentum, small bowel mesentery, ovary) providing additional confirmation.

Prior studies detected MUC2 and MUC5AC in PMP tissue and ascites at the protein level **(5, 58)**. Here, we identified multiple goblet cell-specific secretory products in ascites from PMP patients. More importantly, these proteins were traced to the encoding transcriptome of the accompanying tumor epithelial cells. These tumor cells were present in PMP implants in non-gastrointestinal organs such as the peritoneum and ovary. In addition, manual review of datasets from ovarian and gastric malignant ascites failed to identify these proteins supporting a unique protein profile for appendiceal malignant ascites(59, 60). Altogether, these results for the first time show that mucinous tumor cells with a goblet cell signature contribute to the mucinous phenotype of this cancer.

We identified the trajectory of tumor cell differentiation in both AMN and PMP. This included progenitor states with further differentiation into goblet and metastatic cell states. Notably, PMPs had a specific cell state indicative of their high metastatic potential. We validated the signatures defining these states in PMP tumors from prior studies (3). Importantly, we identified new genes associated with this metastatic cell state including *ANXA1, KLK8* and *FKBP4*.

These genes have been implicated in cancer biology and metastasis among other tumor types but not in appendiceal carcinomas. Further studies of these genes may provide greater insight into the biology of PMP metastasis and their prognostic significance in disease progression.

Our trajectory analysis revealed NOTCH pathway activation in States 4 and 5. A chromosome 9 germline mutation resulting in *NOTCH1* activation has been reported in AMN (17). We identified chromosome 9 amplification in most of the tumors studied here. NOTCH activation has been associated with goblet cell differentiation or hyperplasia in mouse models (38, 61).

These findings suggest an important role for NOTCH signaling in the development of AMN and the first evidence of pathway activation in patient tumors. Current therapy for PMP patients consists of cytoreductive surgery, HIPEC or systemic chemotherapy (2). This study identified activated signaling pathways for potential target therapy including: tetraspanin and kallikrein family genes, MYC, PI3K-AKT-mTOR and RAS signaling pathways. In support, genomic studies have shown, at lower frequency, mutations in *SMAD4, ATM, PIK3CA, AKT* and *JAK3* in AMN and PMP tissues (62). We also identified heterogeneity in the abundance of tumor cell states within the same tumor and at different metastatic sites in the same PMP patient. This may be a source of resistance to treatment regimens.

Both AMN and PMP’s immune TME contained infiltrating T cells. Our results indicated that these cells are similar to those in normal tissue. Additional studies using T-cell receptor (**TCR**) sequencing could further clarify if these cells are bystander cells or have tumor-specificity.

Importantly, these cells retained cytotoxic gene expression suggesting they could be targeted for therapeutic purposes. Recent studies have demonstrated that PMP tumors can respond *ex vivo* to immune checkpoint blockade or neoantigen vaccination (55-57). Further functional characterization of this lymphocyte infiltrate will help clarify the role of these cells in anti-cancer immunity. Strategies that improve cancer cell killing by bystander cells including TCR-independent innate-like killing could also be explored further for these patients (52).

## METHODS

A complete description of methods is provided in the **Supplemental Methods**.

### Samples

Tissues were collected in plain RPMI on ice immediately after resection and dissected with iris scissors. We prepared single cell suspensions using enzymatic and mechanical dissociation (Miltentyi Biotec, Germany) and generated scRNA-seq libraires (10x Genomics, Pleasanton, CA, USA) **(Supplemental Methods)**.

### Batch-corrected integrated analysis and lineage re-clustering

Individual Seurat objects were constructed from each scRNA-seq dataset using Seurat (version 4.0.3) and filtered for low quality cells and computationally identified doublets **(Supplemental Methods)**. Objects were merged and normalized using ‘SCTransform’ (63, 64). To remove batch effects, we integrated all datasets across experimental batches by using a soft variant of k-means clustering implemented in the Harmony algorithm (8) (version 0.1.0). Experimental batch was provided as the grouping variable in the ‘RunHarmony’ function, and this reduction was used in both ‘RunUMAP’ and ‘FindNeighbors’ functions for clustering. First 20 principal components and a resolution of 1 was used for clustering. We used the Adjusted Rand Index (**ARI**) to compare similarity between cluster labels and experimental batch meta data label for each cell. Vector of these respective class labels was supplied to the ‘adjustedRandIndex’ function in mclust package (version 5.4.7) (65).

Following identification of cell lineages based on marker gene expression, we performed a secondary clustering analysis of each lineage. We used the same parameters as described above including integration across experimental batches. From the initial clustering run, we identified clusters that belonged to contaminating cell lineages. These cells were united with their lineage counterparts and a second clustering run was performed yielding final lineage-specific re-clustering results. Following clustering from epithelial cells, low cellularity samples contributing less than 10 total cells were eliminated from downstream analysis.

Data from the ‘RNA’ assay was used for all further downstream analysis with other packages, gene level visualization or differential expression analysis. Data was normalized to the logarithmic scale and effects of variation in sequencing depth were regressed out by including ‘nCount_RNA’ as a parameter in the ‘ScaleData’ function. The ‘DoHeatmap’, ‘FeaturePlot’, ‘DimPlot’, DotPlot’, ‘VlnPlot’ functions were used for visualization.

### Pseudotime trajectory analysis

Expression data obtained from the ‘data’ slot of the ‘RNA’ assay from the Seurat object was used as input to trajectory analysis using Monocle (version 2.20.0) (19) with the ‘importCDS’ function. Top 2000 variable genes identified using variance stabilizing transformation with Seurat function ‘FindVariableFeatures’ were used as input to ‘setorderingfilter’ function that orders cells by examining the pattern of expression of these genes. Next, dimensionality reduction was conducted using the ‘reduceDimension’ function with the DDRTree method and cells were placed onto a pseudotime trajectory using ‘orderCells’. State containing majority of normal cells was set as the root and cells were re-ordered. Resulting states were grouped into 5 major states together with the root state. Differential gene expression between trajectory states was performed using Seurat ‘FindAllMarkers’ function.

### Statistics

Differential gene expression was conducted using the ‘FindAllMarkers’ or ‘FindMarkers’ functions in Seurat using Wilcoxon rank sum test. Parameters provided for these functions were genes detected in at least 25% cells and differential expression threshold of 0.25 log fold change. Significant genes were determined with p-value < 0.05 following Bonferroni correction. Differential pathways and regulatory genes between clusters were determined using ANOVA with post-hoc Tukey Honestly significant difference (HSD). p-value < 0.05 was considered to be significant.

### Study Approval

This study was conducted in compliance with the Helsinki Declaration and approved by the Stanford University School of Medicine Institutional Review Board (IRB-44036). Written informed consent was obtained from all patients.

## DATA AVAILABILITY

Sequencing data will be released under dbGAP identifier phs001818. Cell Ranger count matrices will be available on https://dna-discovery.stanford.edu/research/datasets/.

## AUTHORS’ CONTRIBUTIONS

CA was involved in conception and design of the study, development of methodology, acquisition of data, analysis and interpretation of data and writing of the manuscript. AS was involved in design of the study, development of methodology, analysis and interpretation of data and writing of the manuscript. XB, SG, and JS were involved in the analysis and interpretation of data. GAP and BL were involved in design of the study and clinical translational component. HPJ oversaw the conception and design of the study, data analysis, interpretation of data and writing of the manuscript.

## ACKNOWLEDGEMENTS

We are grateful to all patients who participated in the study. This work was supported by the Don and Ruth Seiler Fund. Other support came from US National Institutes of Health grants R01HG006137 (HPJ) and U01CA217875 (HPJ, AS). HPJ also received support from the Clayville Foundation. Figures 1A, 5A and graphical abstract were created using BioRender.com.

## SUPPLEMENTAL DATA

Supplemental Methods. Format: PDF

Supplemental Figure S1-S4. Format: PDF

Supplemental Tables S1 – S7. Format: XLSX

---

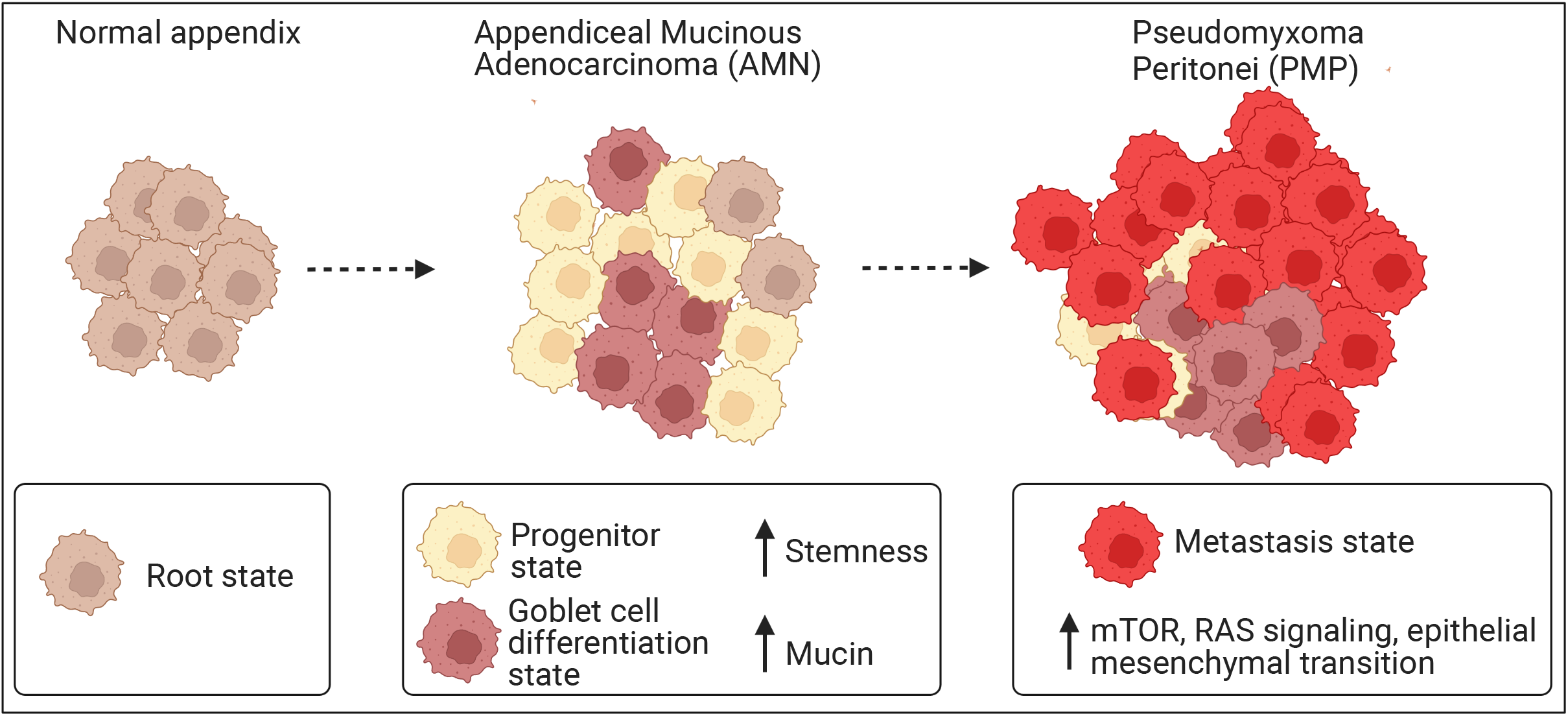

## REFERENCES

1. Choudry HA, Pai RK. Management of Mucinous Appendiceal Tumors. Ann Surg Oncol. 2018;25(8):2135–44.

2. Bartlett DJ, Thacker PG, Jr., Grotz TE, Graham RP, Fletcher JG, VanBuren WM, et al. Mucinous appendiceal neoplasms: classification, imaging, and HIPEC. Abdom Radiol (NY). 2019;44(5):1686–702.

3. Levine EA, Votanopoulos KI, Qasem SA, Philip J, Cummins KA, Chou JW, et al. Prognostic Molecular Subtypes of Low-Grade Cancer of the Appendix. J Am Coll Surg. 2016;222(4):493–503.

4. O’Connell JT, Hacker CM, Barsky SH. MUC2 is a molecular marker for pseudomyxoma peritonei. Mod Pathol. 2002;15(9):958–72.

5. O’Connell JT, Tomlinson JS, Roberts AA, McGonigle KF, Barsky SH. Pseudomyxoma peritonei is a disease of MUC2-expressing goblet cells. Am J Pathol. 2002;161(2):551–64.

6. Carr NJ, Bibeau F, Bradley RF, Dartigues P, Feakins RM, Geisinger KR, et al. The histopathological classification, diagnosis and differential diagnosis of mucinous appendiceal neoplasms, appendiceal adenocarcinomas and pseudomyxoma peritonei. Histopathology. 2017;71(6):847–58.

7. Levine EA, Blazer DG, 3rd, Kim MK, Shen P, Stewart JHt, Guy C, et al. Gene expression profiling of peritoneal metastases from appendiceal and colon cancer demonstrates unique biologic signatures and predicts patient outcomes. J Am Coll Surg. 2012;214(4):599-606; discussion −7.

8. Korsunsky I, Millard N, Fan J, Slowikowski K, Zhang F, Wei K, et al. Fast, sensitive and accurate integration of single-cell data with Harmony. Nat Methods. 2019;16(12):1289–96.

9. Hubert L, Arabie P. Comparing partitions. Journal of Classification. 1985;2(1):193–218.

10. Elmentaite R, Kumasaka N, Roberts K, Fleming A, Dann E, King HW, et al. Cells of the human intestinal tract mapped across space and time. Nature. 2021;597(7875):250–5.

11. Aran D, Looney AP, Liu L, Wu E, Fong V, Hsu A, et al. Reference-based analysis of lung single-cell sequencing reveals a transitional profibrotic macrophage. Nat Immunol. 2019;20(2):163–72.

12. Ramachandran P, Dobie R, Wilson-Kanamori JR, Dora EF, Henderson BEP, Luu NT, et al. Resolving the fibrotic niche of human liver cirrhosis at single-cell level. Nature. 2019;575(7783):512–8.

13. Haber AL, Biton M, Rogel N, Herbst RH, Shekhar K, Smillie C, et al. A single-cell survey of the small intestinal epithelium. Nature. 2017;551(7680):333–9.

14. Nguyen B, Sanchez-Vega F, Fong CJ, Chatila WK, Boroujeni AM, Pareja F, et al. The genomic landscape of carcinomas with mucinous differentiation. Sci Rep. 2021;11(1):9478.

15. Patel AP, Tirosh I, Trombetta JJ, Shalek AK, Gillespie SM, Wakimoto H, et al. Single-cell RNA-seq highlights intratumoral heterogeneity in primary glioblastoma. Science. 2014;344(6190):1396–401.

16. Tirosh I, Izar B, Prakadan SM, Wadsworth MH, 2nd, Treacy D, Trombetta JJ, et al. Dissecting the multicellular ecosystem of metastatic melanoma by single-cell RNA-seq. Science. 2016;352(6282):189–96.

17. LaFramboise WA, Pai RK, Petrosko P, Belsky MA, Dhir A, Howard PG, et al. Discrimination of low- and high-grade appendiceal mucinous neoplasms by targeted sequencing of cancer-related variants. Mod Pathol. 2019;32(8):1197–209.

18. Alakus H, Babicky ML, Ghosh P, Yost S, Jepsen K, Dai Y, et al. Genome-wide mutational landscape of mucinous carcinomatosis peritonei of appendiceal origin. Genome Med. 2014;6(5):43.

19. Qiu X, Mao Q, Tang Y, Wang L, Chawla R, Pliner HA, et al. Reversed graph embedding resolves complex single-cell trajectories. Nat Methods. 2017;14(10):979–82.

20. Moreno CS. SOX4: The unappreciated oncogene. Semin Cancer Biol. 2020;67(Pt 1):57–64.

21. Wang H, Yu Z, Huo S, Chen Z, Ou Z, Mai J, et al. Overexpression of ELF3 facilitates cell growth and metastasis through PI3K/Akt and ERK signaling pathways in non-small cell lung cancer. Int J Biochem Cell Biol. 2018;94:98–106.

22. Gracz AD, Samsa LA, Fordham MJ, Trotier DC, Zwarycz B, Lo YH, et al. Sox4 Promotes Atoh1-Independent Intestinal Secretory Differentiation Toward Tuft and Enteroendocrine Fates. Gastroenterology. 2018;155(5):1508–23 e10.

23. Ng AY, Waring P, Ristevski S, Wang C, Wilson T, Pritchard M, et al. Inactivation of the transcription factor Elf3 in mice results in dysmorphogenesis and altered differentiation of intestinal epithelium. Gastroenterology. 2002;122(5):1455–66.

24. Noah TK, Kazanjian A, Whitsett J, Shroyer NF. SAM pointed domain ETS factor (SPDEF) regulates terminal differentiation and maturation of intestinal goblet cells. Exp Cell Res. 2010;316(3):452–65.

25. Heo K, Lee S. TSPAN8 as a Novel Emerging Therapeutic Target in Cancer for Monoclonal Antibody Therapy. Biomolecules. 2020;10(3).

26. Zhang J, Zhu Z, Miao Z, Huang X, Sun Z, Xu H, et al. The Clinical Significance and Mechanisms of REG4 in Human Cancers. Front Oncol. 2020;10:559230.

27. Chai CY, Zhang Y, Song J, Lin SC, Sun S, Chang IW. VNN1 overexpression is associated with poor response to preoperative chemoradiotherapy and adverse prognosis in patients with rectal cancers. Am J Transl Res. 2016;8(10):4455–63.

28. Liu GJ, Wang YJ, Yue M, Zhao LM, Guo YD, Liu YP, et al. High expression of TCN1 is a negative prognostic biomarker and can predict neoadjuvant chemosensitivity of colon cancer. Sci Rep. 2020;10(1):11951.

29. Nugteren S, Samsom JN. Secretory Leukocyte Protease Inhibitor (SLPI) in mucosal tissues: Protects against inflammation, but promotes cancer. Cytokine Growth Factor Rev. 2021;59:22–35.

30. Santiago-Sanchez GS, Pita-Grisanti V, Quinones-Diaz B, Gumpper K, Cruz-Monserrate Z, Vivas-Mejia PE. Biological Functions and Therapeutic Potential of Lipocalin 2 in Cancer. Int J Mol Sci. 2020;21(12).

31. Shvartsur A, Bonavida B. Trop2 and its overexpression in cancers: regulation and clinical/therapeutic implications. Genes Cancer. 2015;6(3-4):84–105.

32. Gebauer F, Wicklein D, Horst J, Sundermann P, Maar H, Streichert T, et al. Carcinoembryonic antigen-related cell adhesion molecules (CEACAM) 1, 5 and 6 as biomarkers in pancreatic cancer. PLoS One. 2014;9(11):e113023.

33. Araujo TG, Mota STS, Ferreira HSV, Ribeiro MA, Goulart LR, Vecchi L. Annexin A1 as a Regulator of Immune Response in Cancer. Cells. 2021;10(9).

34. Hua Q, Li T, Liu Y, Shen X, Zhu X, Xu P. Upregulation of KLK8 Predicts Poor Prognosis in Pancreatic Cancer. Front Oncol. 2021;11:624837.

35. Xiong H, Chen Z, Zheng W, Sun J, Fu Q, Teng R, et al. FKBP4 is a malignant indicator in luminal A subtype of breast cancer. J Cancer. 2020;11(7):1727–36.

36. Hanzelmann S, Castelo R, Guinney J. GSVA: gene set variation analysis for microarray and RNA-seq data. BMC Bioinformatics. 2013;14:7.

37. Liberzon A, Birger C, Thorvaldsdottir H, Ghandi M, Mesirov JP, Tamayo P. The Molecular Signatures Database (MSigDB) hallmark gene set collection. Cell Syst. 2015;1(6):417–25.

38. Feng Y, Bommer GT, Zhao J, Green M, Sands E, Zhai Y, et al. Mutant KRAS promotes hyperplasia and alters differentiation in the colon epithelium but does not expand the presumptive stem cell pool. Gastroenterology. 2011;141(3):1003-13 e1-10.

39. Aibar S, Gonzalez-Blas CB, Moerman T, Huynh-Thu VA, Imrichova H, Hulselmans G, et al. SCENIC: single-cell regulatory network inference and clustering. Nat Methods. 2017;14(11):1083–6.

40. Treveil A, Sudhakar P, Matthews ZJ, Wrzesinski T, Jones EJ, Brooks J, et al. Regulatory network analysis of Paneth cell and goblet cell enriched gut organoids using transcriptomics approaches. Mol Omics. 2020;16(1):39–58.

41. Hickey JW, Becker WR, Nevins SA, Horning A, Perez AE, Chiu R, et al. High Resolution Single Cell Maps Reveals Distinct Cell Organization and Function Across Different Regions of the Human Intestine. bioRxiv. 2021:2021.11.25.469203.

42. Xiong Y, Feng Y, Zhao J, Lei J, Qiao T, Zhou Y, et al. TFAP2A potentiates lung adenocarcinoma metastasis by a novel miR-16 family/TFAP2A/PSG9/TGF-beta signaling pathway. Cell Death Dis. 2021;12(4):352.

43. Liu Y, Wang G, Yang Y, Mei Z, Liang Z, Cui A, et al. Increased TEAD4 expression and nuclear localization in colorectal cancer promote epithelial-mesenchymal transition and metastasis in a YAP-independent manner. Oncogene. 2016;35(21):2789–800.

44. Geller A, Yan J. The Role of Membrane Bound Complement Regulatory Proteins in Tumor Development and Cancer Immunotherapy. Front Immunol. 2019;10:1074.

45. Lin YL, Ma R, Li Y. The biological basis and function of GNAS mutation in pseudomyxoma peritonei: a review. J Cancer Res Clin Oncol. 2020;146(9):2179–88.

46. Bern M, Kil YJ, Becker C. Byonic: Advanced Peptide and Protein Identification Software. Current Protocols in Bioinformatics. 2012;40(1):13.20.1-13.20.14.

47. Elias JE, Gygi SP. Target-decoy search strategy for increased confidence in large-scale protein identifications by mass spectrometry. Nat Methods. 2007;4(3):207–14.

48. Kim YS, Ho SB. Intestinal goblet cells and mucins in health and disease: recent insights and progress. Curr Gastroenterol Rep. 2010;12(5):319–30.

49. Nieto P, Elosua-Bayes M, Trincado JL, Marchese D, Massoni-Badosa R, Salvany M, et al. A single-cell tumor immune atlas for precision oncology. Genome Res. 2021;31(10):1913–26.

50. van der Leun AM, Thommen DS, Schumacher TN. CD8(+) T cell states in human cancer: insights from single-cell analysis. Nat Rev Cancer. 2020;20(4):218–32.

51. Su J, Jin G, Votanopoulos KI, Craddock L, Shen P, Chou JW, et al. Prognostic Molecular Classification of Appendiceal Mucinous Neoplasms Treated with Cytoreductive Surgery and Hyperthermic Intraperitoneal Chemotherapy. Ann Surg Oncol. 2020;27(5):1439–47.

52. Meier SL, Satpathy AT, Wells DK. Bystander T cells in cancer immunology and therapy. Nat Cancer. 2022;3(2):143–55.

53. Zheng C, Zheng L, Yoo JK, Guo H, Zhang Y, Guo X, et al. Landscape of Infiltrating T Cells in Liver Cancer Revealed by Single-Cell Sequencing. Cell. 2017;169(7):1342–56 e16.

54. Guo X, Zhang Y, Zheng L, Zheng C, Song J, Zhang Q, et al. Global characterization of T cells in non-small-cell lung cancer by single-cell sequencing. Nat Med. 2018;24(7):978–85.

55. Forsythe SD, Erali RA, Sasikumar S, Laney P, Shelkey E, D’Agostino R, Jr., et al. Organoid Platform in Preclinical Investigation of Personalized Immunotherapy Efficacy in Appendiceal Cancer: Feasibility Study. Clin Cancer Res. 2021;27(18):5141–50.

56. Flatmark K, Torgunrud A, Fleten KG, Davidson B, Juul HV, Mensali N, et al. Peptide vaccine targeting mutated GNAS: a potential novel treatment for pseudomyxoma peritonei. J Immunother Cancer. 2021;9(10).

57. Weitz J, Hurtado de Mendoza T, Tiriac H, Lee J, Sun S, Garg B, et al. An Ex Vivo Organotypic Culture Platform for Functional Interrogation of Human Appendiceal Cancer Reveals a Prominent and Heterogenous Immunological Landscape. Clin Cancer Res. 2022;28(21):4793–806.

58. Mall AS, Chirwa N, Govender D, Lotz Z, Tyler M, Rodrigues J, et al. MUC2, MUC5AC and MUC5B in the mucus of a patient with pseudomyxoma peritonei: biochemical and immunohistochemical study. Pathol Int. 2007;57(8):537–47.

59. Jin J, Son M, Kim H, Kim H, Kong SH, Kim HK, et al. Comparative proteomic analysis of human malignant ascitic fluids for the development of gastric cancer biomarkers. Clin Biochem. 2018;56:55–61.

60. Shender VO, Pavlyukov MS, Ziganshin RH, Arapidi GP, Kovalchuk SI, Anikanov NA, et al. Proteome-metabolome profiling of ovarian cancer ascites reveals novel components involved in intercellular communication. Mol Cell Proteomics. 2014;13(12):3558–71.

61. Zecchini V, Domaschenz R, Winton D, Jones P. Notch signaling regulates the differentiation of post-mitotic intestinal epithelial cells. Genes Dev. 2005;19(14):1686–91.

62. Stein A, Strong E, Clark Gamblin T, Clarke C, Tsai S, Thomas J, et al. Molecular and Genetic Markers in Appendiceal Mucinous Tumors: A Systematic Review. Ann Surg Oncol. 2020;27(1):85–97.

63. Butler A, Hoffman P, Smibert P, Papalexi E, Satija R. Integrating single-cell transcriptomic data across different conditions, technologies, and species. Nat Biotechnol. 2018;36(5):411–20.

64. Hafemeister C, Satija R. Normalization and variance stabilization of single-cell RNA-seq data using regularized negative binomial regression. Genome Biol. 2019;20(1):296.

65. Scrucca L, Fop M, Murphy TB, Raftery AE. mclust 5: Clustering, Classification and Density Estimation Using Gaussian Finite Mixture Models. The R Journal. 2016;8:289--317.

