## Supplemental materials for "Distinct cell states define the developmental trajectories of mucinous appendiceal neoplasms towards pseudomyxoma metastases"

**TITLE**

**INSTITUTIONS**

[1] Division of Surgical Oncology, Department of Surgery, Stanford University, Stanford, CA 94305, United States

[2] Division of Oncology, Department of Medicine, Stanford University School of Medicine, Stanford, CA, United States

[3] Department of Pathology, Stanford University School of Medicine, Stanford, CA USA

**CORRESPONDING AUTHOR**

Hanlee P. Ji

Mailing address: CCSR 2245, 269 Campus Drive, Stanford, CA-94305, USA

### SUPPLEMENTAL METHODS

#### Tissue and ascites processing

Dissected tissue was subjected to enzymatic and mechanical dissociation using human tumor dissociation kit (Miltentyi Biotec, Germany) with the Gentle MACS Octodissociator (Miltenyi) as per manufacturer's protocol with the '37_h_TDK_3' program. Dissociated cells were incubated with RBC lysis buffer (155 mM ammonium chloride, 10 mM potassium bicarbonate, 0.1 mM EDTA) for 5 minutes followed by neutralization with double volume PBS. All centrifugation steps were carried out at 500g for 5 minutes. Cells were cryo-frozen in freezing media consisting of 10% DMSO in 90% FBS (Thermo Fisher Scientific, Waltham, MA) in a CoolCell freezing container (Larkspur, CA) at -80ºC for 48 hours followed by storage in liquid nitrogen.

At time of use, cryo-frozen cells were rapidly thawed in a bead bath at 37ºC. Next, dissociated cells were washed twice in RPMI + 10% FBS. Cells were subsequently filtered through a 70 µm followed by 40 µm filter (Corning Sterile Cell Strainer, Fischer-Scientific). Live cell counts were obtained on a Countess 3 Automated Cell Counter (Thermo Fisher Scientific) using 1:1 trypan blue dilution. Cells were concentrated between 100-1500 live cells/µl and used for loading single-cell reactions.

Ascites was collected fresh in plastic buckets upon entry into the abdomen. Ascites was kept at 37ºC to avoid gel formation or solidification. Volume to be used was subsequently processed by sequential filtration through 100, 70 and 40 µm strainers. Ascites was preserved in freezing media following RBC lysis as described above.

**Single cell RNA-sequencing (scRNA-seq) library preparation**

Samples from each patient were processed in one batch for library preparation. Chromium Single Cell 5′ Library & Gel Bead Kit v1.0 or v1.1 (10x Genomics, Pleasanton, CA, USA) were used as per manufacturer's protocol. For a given sample, 10000 cells were targeted for tissue dissociation suspensions with 18 PCR cycles for cDNA amplification. 14 or 16 PCR cycles were used for library amplification when amplified cDNA input was between 25-50 ng or less than 25 ng respectively. Libraries were sequenced on Illumina NovaSeq or NextSeq platforms.

**scRNA-seq data processing for individual samples**

Cell Ranger (10x Genomics) version 3.1.0 'mkfastq' and 'count' commands were used with default parameters and alignment to GRCh38 to generate matrix of unique molecular identifier **(UMI)** counts per gene and associated cell barcode. We constructed Seurat objects from each dataset using Seurat (version 3.0.2) (1, 2) to apply quality control filters. We removed cells that expressed fewer than 200 genes, had greater than 30% mitochondrial genes or had UMI counts greater than 5000 indicatives of potential doublets. We removed genes that were detected in less than 3 cells. We normalized data using 'SCTransform' and used first 50 principal components with a resolution of 0.8 for clustering. We then removed computationally identified doublets from each dataset using DoubletFinder (version 2.0.2) (3). The 'pN' value was set to default value of 0.25 as the proportion of artificial doublets. 'nExP' was set to expected doublet rate according to Chromium Single Cell 3' v2 reagents kits user guide (10x Genomics). These parameters were used as input to the 'doubletFinder_v3' function with number of principal components set to 20 to identify doublet cells.

### Copy number analysis

InferCNV ( version 1.2.3) (4) was used to infer large-scale copy number variations in tumor epithelial cells identified as goblet cells following reference transcriptome mapping. As a reference control, we used all epithelial cells from normal appendiceal tissue and 500 cells each from all myeloid, lymphocyte and stromal cells using random sampling with seed set to 123. Count data was used as input. Filtering, normalization and centering by normal gene expression was performed using default parameters. A cut-off of 0.1 was used for the minimum average read counts per gene among reference cells. An additional denoising filter was used with a threshold of 0.2. Copy number variation was predicted using the default six state Hidden Markov Model.

### Cell annotation using reference atlases

Gut epithelium single-cell reference atlas (5) was downloaded from https://www.gutcellatlas.org/#datasets. zellkonverter (version 1.2.1) (6) was used to convert data into a Seurat object. Tumor infiltrating immune cell atlas (7) Seurat object and metadata was obtained from 10.5281/zenodo.4263972. The atlas was filtered for cells belonging to T and NK cell lineages. All counts from reference atlases were normalized to the logarithmic scale and used as a reference for automated annotation per cell using SingleR (version 1.14.1) (8). Raw counts were used to annotate test datasets. Labels were predicted for each cell in the test dataset using Spearman correlation with Wilcoxon Rank Sum test for marker gene detection in the reference dataset. This was performed using the function ‘SingleR’ with default parameters.

Following automated label assignment using this method, we confirmed results by examining marker gene expression. Cell labels were then reannotated in case of misassignment in keeping with current recommended best practices (9). Granular labels from the gut cell atlas were combined for downstream analysis. Goblet, BEST2+ goblet and microfold cells were regrouped together as goblet cells. Colonocyte, BEST4+ epithelial, enterocyte, transit amplifying cells and a single detected Paneth cell were regrouped as enterocytes. Stem, distal progenitor and CLDN10+ cells were regrouped together as stem cells. Enteroendocrine D, enteroendocrine NPW+ and enteroendocrine TAC+ cells were regrouped together as enteroendocrine cells.

T helper cells from tumor immune cell atlas were renamed as 'TFh-like' cells. Effector memory CD8 T cells, Cytotoxic CD8 T cells and Proliferative T cells were grouped together as 'Cytotoxic_CD8' based on marker gene expression. Th17 cells, Transitional memory CD4 T cells, Recently activated CD4 T cells, and Naive-memory CD4 T cells were grouped together as 'Naïve-like' based on marker gene expression.

**Cell type/state abundance analysis**

We used a generalized linear regression model to determine if the abundance of cell types/states corelates with condition by modelling the detected number of cells in each cell type/state as a random variable using a Poisson process as previously described (10, 11). Log transformed cell numbers were used as an offset variable. Condition (Normal, AMN or PMP) for each sample was fitted as the covariate in 'glm' function of the 'stats' package (version 4.1.0). P-value for significance of effect of condition was obtained using Wald test on regression coefficient.

**Pathway analysis**

Hallmark gene sets were downloaded from MSigDB version 6.2 (12, 13) and read using 'getGmt' function in GSEAbase (version 1.54.0) (14). Enrichment scores for each cell were calculated with GSVA (version 1.40.1) package using function 'gsva' and parameters 'kcdf' and 'mx.diff' set to Gaussian and true respectively. Expression matrix derived from the data slot of the RNA assay in the Seurat object was used as input. Average enrichment score was determined for each trajectory branch. Significantly different scores across clusters were determined by ANOVA with post-hoc Tukey Honestly significant difference (HSD) corrected p-value < 0.05 and represented as heatmap with Ward.D2 clustering.

We used the 'AddModuleScore' function in Seurat to calculate the average expression of gene signatures. This included gel-forming mucins (*MUC2*, *MUC5B*, *MUC5A*, *MUC6*, *MUC19*) (15), CD8 cytotoxicity (16, 17) and CD8 exhaustion (18) (**Supplemental Table 7**). Using this function, genes of interest were first binned into 24 bins of expression levels based on their average expression. From each bin, control genes were randomly selected using default parameters used in this function. Finally, average expression score was calculated as the difference between average expression of gene set of interest and average expression of control genes. Expression between clusters was compared using ANOVA with post-hoc Tukey Honestly significant difference (HSD).

**Regulon analysis**

Gene regulatory networks were constructed for each cell using SCENIC (version 1.1.1.5) (19) with dependencies AUCell (version 1.5.5), Rcistarget (version 1.3.4) and Genie3 (version 1.5.4). Average AUC scores were calculated per cluster and compared using ANOVA with post-hoc

Tukey Honestly significant difference (HSD) and represented as heatmap with Ward.D2 clustering.

**Bulk gene expression datasets and signature enrichment analysis**

Normalized bulk gene expression data was obtained from Gene Expression Omnibus series GSE75535 with platforms GPL13667 and GPL571 (20), using the 'getGEO' function from the GEOquery R package (version 2.60.0). This data set was generated from appendiceal tumors where the total RNA was profiled on Affymetrix U133A GeneChips and Affymetrix U219 GeneAtlas Array Strips, respectively. Average counts were used for probes matching duplicate gene names. GSVA was conducted using parameters described above. Gene sets for progenitor, goblet and metastasis states were constructed using the results of differential expression of trajectory analysis states from scRNA-seq data. Marker genes were uniquely assigned to the cell state with higher fold change value. Only genes that were present in the microarray datasets were included. Spearman correlation between enrichment scores was obtained using 'cor.test' function from the stats R package (version 4.1.0).

**Histopathology**

A portion of surgically resected tissue was fixed in 10% formalin for 24 hours at room temperature. Paraffin embedding and processing with hematoxylin and eosin staining was conducted by the Human Pathology Histology Services core facility at Stanford University. Slides were reviewed by a board-certified pathologist. We additionally reviewed clinical histopathology reports for all patients.

### Diagnostic Cancer Gene Sequencing

Three patients (P5853, P8605, P8629) underwent diagnostic cancer gene sequencing as part of their clinical care. We reviewed these reports for mutation status in *KRAS* and *GNAS* genes.

### Immunohistochemistry

Immunohistochemistry (IHC) staining was performed for MUC2 (#sc-7314, Santa Cruz Biotechnology, TX, USA, dilution 1:250), AGR2 (#HPA007912, Sigma-Aldrich, Inc., MO, USA, dilution 1:500) and Annexin A1 (#32934, Cell Signaling Technology, MA, USA, dilution 1:400) using manufacturer's recommended protocol. SignalStain Citrate Unmasking Solution and Animal-Free Blocking Solution (Cell Signaling Technology) were used for antigen retrieval and blocking respectively. Species-specific isotype antibody (Cell Signaling Technology) at the same concentration as primary antibody was used as isotype staining control. Whole slide images were obtained using Aperio AT2 whole slide scanner (Leica Biosystems Inc., IL, USA) and QuPath (version 0.2.3) software (21).

### Mass spectrometry of mucinous ascites

Protein samples were precipitated with 4X volume of acetone and incubated at -80°C overnight. Air-dried protein pellets were reconstituted in 200ul of 50mM ammonium bicarbonate buffer, sonicated, and vortexed to solubilize proteins. The samples were then reduced with 10mM DTT at 55ºC for 30 minutes followed by alkylation with 30mM acrylamide for 30 minutes at room temperature. 1µg of Trypsin/LysC protease (Promega) was added to each sample for digestion at 37ºC overnight. After digestion, the reaction was quenched using 1% formic acid and peptides were de-salted on C18 Monospin reversed phase columns (GL Sciences). Peptide quantification was performed with the Pierce Quantitative Fluorometric Peptide Assay kit (Thermo Fisher Scientific). The peptide mixture was dried by speed vac before dissolution in reconstitution buffer (2% acetonitrile with 0.1% formic acid). 0.5µg was used for subsequent LC-MS/MS analysis.

Mass spectrometry experiments were performed on a Q Exactive HF-X Hybrid Quadrupole - Orbitrap mass spectrometer (Thermo Scientific, San Jose, CA) with liquid chromatography using a Nanoacquity UPLC (Waters Corporation, Milford, MA). For a typical LCMS experiment, a flow rate of 450 nL/min is used, where mobile phase A was 0.2% formic acid in water and mobile phase B was 0.2% formic acid in acetonitrile. For this analysis, a 50cm µPAC column (Pharmafluidics) was used. Peptides were directly injected onto the analytical column using a gradient (2-45% B, followed by a high-B wash) of 80min. The mass spectrometer was operated in a data dependent fashion using HCD fragmentation for MS/MS spectra generation.

For data analysis, the .RAW data files were processed using Byonic (Protein Metrics, Cupertino, CA) to identify peptides and infer proteins (22). The readout from the data processing is a protein list ranked by the base-10 logarithm of the protein p-value. The p-value represents the likelihood of the peptide spectrum match being random. The confidence threshold for an accurate call requires a protein with log p-value at least 2.0 fold lower than the log p-value of the top decoy protein.

Proteolysis with Trypsin/LysC was assumed to be semi-specific allowing for N-ragged cleavage with up to two missed cleavage sites. Precursor and fragment mass accuracies were held within 12 ppm. Cysteine modified with propionamide were set as fixed modifications in the search. Proteins were held to a false discovery rate of 1%, using standard reverse-decoy technique (23). Protein names were mapped to gene names using https://www.uniprot.org/uploadlists/. We identified significantly enriched proteins with logarithmic p-values two fold lower than those of decoy proteins (22).

Additional analysis or visualization was conducted using R packages tidyverse (version 1.3.1), dplyr (version 1.0.6), ggplot2 (version 3.3.3), broom (version 0.7.6), pheatmap (version 1.0.12) and viridis (version 0.6.1) in R version 4.1.0.

### REFERENCES


1. Butler A, Hoffman P, Smibert P, Papalexi E, Satija R. Integrating single-cell transcriptomic data across different conditions, technologies, and species. Nat Biotechnol. 2018;36(5):411-20.

2. Hafemeister C, Satija R. Normalization and variance stabilization of single-cell RNA-seq data using regularized negative binomial regression. Genome Biol. 2019;20(1):296.

3. McGinnis CS, Murrow LM, Gartner ZJ. DoubletFinder: Doublet Detection in Single-Cell RNA Sequencing Data Using Artificial Nearest Neighbors. Cell Syst. 2019;8(4):329-37 e4.

4. Tickle T, Tirosh I, Georgescu C, Brown M, Haas B. inferCNV of the Trinity CTAT Project: Klarman Cell Observatory, Broad Institute of MIT and Harvard, Cambridge, MA, USA; 2019 [Available from: https://github.com/broadinstitute/inferCNV.

5. Elmentaite R, Kumasaka N, Roberts K, Fleming A, Dann E, King HW, et al. Cells of the human intestinal tract mapped across space and time. Nature. 2021;597(7875):250-5.

6. Zappia L, Lun A. zellkonverter: Conversion Between scRNA-seq Objects. R package version 1.5.0, https://github.com/theislab/zellkonverter. 2021.

7. Nieto P, Elosua-Bayes M, Trincado JL, Marchese D, Massoni-Badosa R, Salvany M, et al. A single-cell tumor immune atlas for precision oncology. Genome Res. 2021;31(10):1913-26.

8. Aran D, Looney AP, Liu L, Wu E, Fong V, Hsu A, et al. Reference-based analysis of lung single-cell sequencing reveals a transitional profibrotic macrophage. Nat Immunol. 2019;20(2):163-72.

9. Clarke ZA, Andrews TS, Atif J, Pouyabahar D, Innes BT, MacParland SA, et al. Tutorial: guidelines for annotating single-cell transcriptomic maps using automated and manual methods. Nat Protoc. 2021;16(6):2749-64.

10. Ramachandran P, Dobie R, Wilson-Kanamori JR, Dora EF, Henderson BEP, Luu NT, et al. Resolving the fibrotic niche of human liver cirrhosis at single-cell level. Nature. 2019;575(7783):512-8.

11. Haber AL, Biton M, Rogel N, Herbst RH, Shekhar K, Smillie C, et al. A single-cell survey of the small intestinal epithelium. Nature. 2017;551(7680):333-9.

12. Hanzelmann S, Castelo R, Guinney J. GSVA: gene set variation analysis for microarray and RNA-seq data. BMC Bioinformatics. 2013;14:7.

13. Liberzon A, Birger C, Thorvaldsdottir H, Ghandi M, Mesirov JP, Tamayo P. The Molecular Signatures Database (MSigDB) hallmark gene set collection. Cell Syst. 2015;1(6):417-25.

14. Morgan M, Falcon S, Gentleman R. GSEABase: Gene set enrichment data structures and methods. R package version 1.46.0. 2019.

15. Nguyen B, Sanchez-Vega F, Fong CJ, Chatila WK, Boroujeni AM, Pareja F, et al. The genomic landscape of carcinomas with mucinous differentiation. Sci Rep. 2021;11(1):9478.

16. Tirosh I, Izar B, Prakadan SM, Wadsworth MH, 2nd, Treacy D, Trombetta JJ, et al. Dissecting the multicellular ecosystem of metastatic melanoma by single-cell RNA-seq. Science. 2016;352(6282):189-96.

17. Guo X, Zhang Y, Zheng L, Zheng C, Song J, Zhang Q, et al. Global characterization of T cells in non-small-cell lung cancer by single-cell sequencing. Nat Med. 2018;24(7):978-85.

18. Zheng C, Zheng L, Yoo JK, Guo H, Zhang Y, Guo X, et al. Landscape of Infiltrating T Cells in Liver Cancer Revealed by Single-Cell Sequencing. Cell. 2017;169(7):1342-56 e16.

19. Aibar S, Gonzalez-Blas CB, Moerman T, Huynh-Thu VA, Imrichova H, Hulselmans G, et al. SCENIC: single-cell regulatory network inference and clustering. Nat Methods. 2017;14(11):1083-6.

20. Levine EA, Votanopoulos KI, Qasem SA, Philip J, Cummins KA, Chou JW, et al. Prognostic Molecular Subtypes of Low-Grade Cancer of the Appendix. J Am Coll Surg. 2016;222(4):493-503.

21. Bankhead P, Loughrey MB, Fernandez JA, Dombrowski Y, McArt DG, Dunne PD, et al. QuPath: Open source software for digital pathology image analysis. Sci Rep. 2017;7(1):16878.

22. Bern M, Kil YJ, Becker C. Byonic: Advanced Peptide and Protein Identification Software. Current Protocols in Bioinformatics. 2012;40(1):13.20.1-13.20.14.

23. Elias JE, Gygi SP. Target-decoy search strategy for increased confidence in large-scale protein identifications by mass spectrometry. Nat Methods. 2007;4(3):207-14.

### LEGENDS TO SUPPLEMENTAL FIGURES

**Supplemental Figure 1.** (A) Representative images of hematoxylin and eosin staining of respective tumor samples. Scale bar = 100 μm. LAMN: low-grade appendiceal mucinous neoplasm, LMCP: low-grade mucinous carcinoma peritonei, HMCP: high-grade mucinous carcinoma peritonei. (B) UMAP representation of dimensionally reduced data following batch correction and graph-based clustering of epithelial cells from all datasets annotated by condition. (C-D) Violin plots depicting expression of (C) *MUC2* gene or (D) gel-forming Mucins gene set with ANOVA Tukey HSD p-value in normal, AMN and PMP epithelial cells.

**Supplemental Figure 2.** Heatmap representation of inferred single-cell CNV profiles of goblet tumor epithelial cells per respective sample compared to reference cells from normal epithelium, immune and stromal cells. CNV legend: state 1 = complete loss, state 2 = loss of one copy, state 3 = neutral, state 4 = addition of one copy, state 5 = addition of two copies, state 6 = addition of more than 2 copies.

**Supplemental Figure 3.** Heatmap depicting average expression levels of respective genes per epithelial cell state per patient for (A) *ANXA2* and (B) *LCN2*. The '*' indicates significant differential expression (adjusted p-value < 0.05) for that cell state compared to the other cell states in respective patient.

**Supplemental Figure 4**. (A-C) Violin plots indicating expression of selected differentially expressed genes in tumor epithelial cells per PMP lesion site in respective patients. The '*' indicates a lesion with significant differential expression (adjusted p-value < 0.05) for that gene compared to the other lesions.

### Supplemental Figure 1

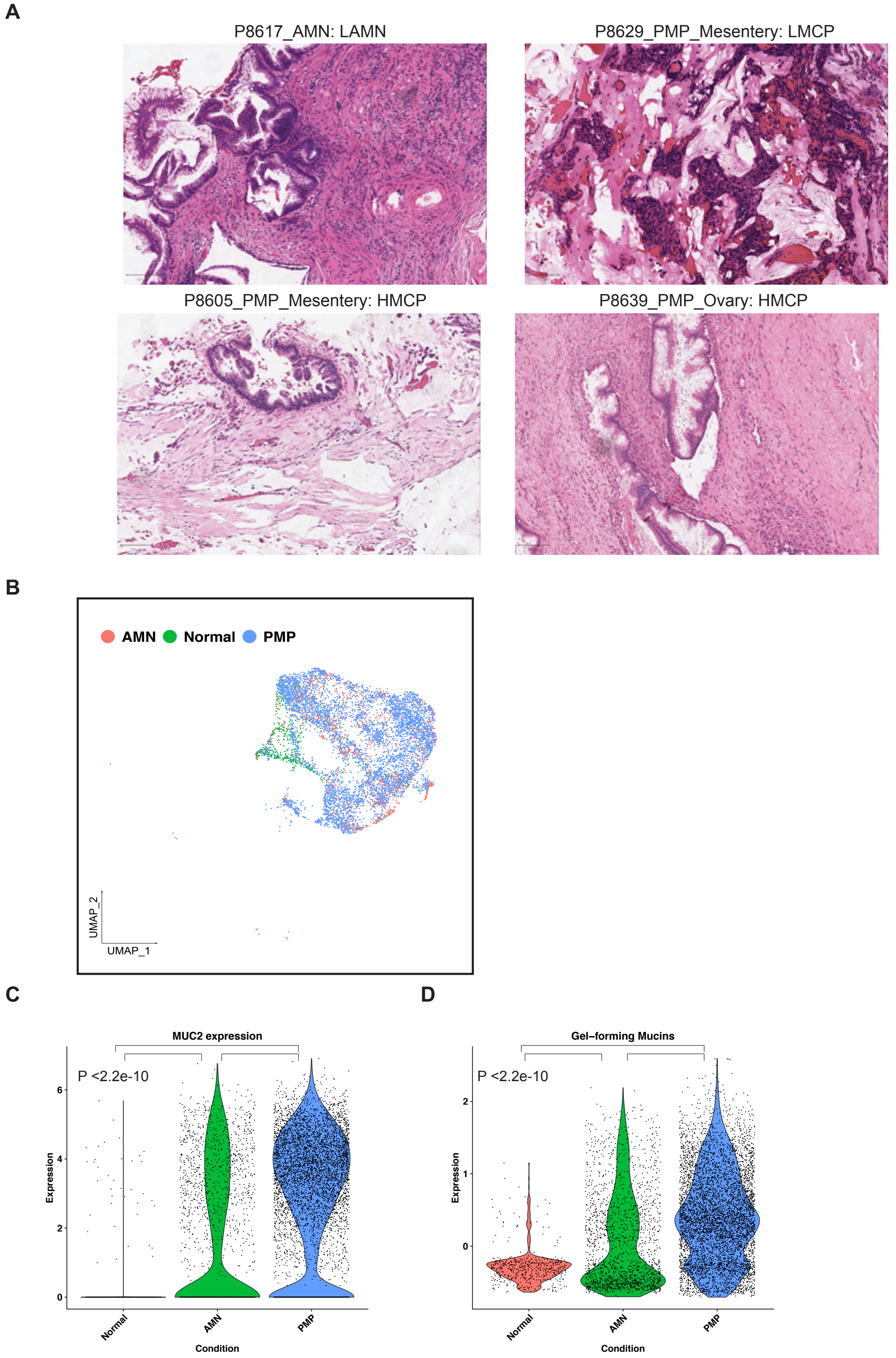

### Supplemental Figure 2

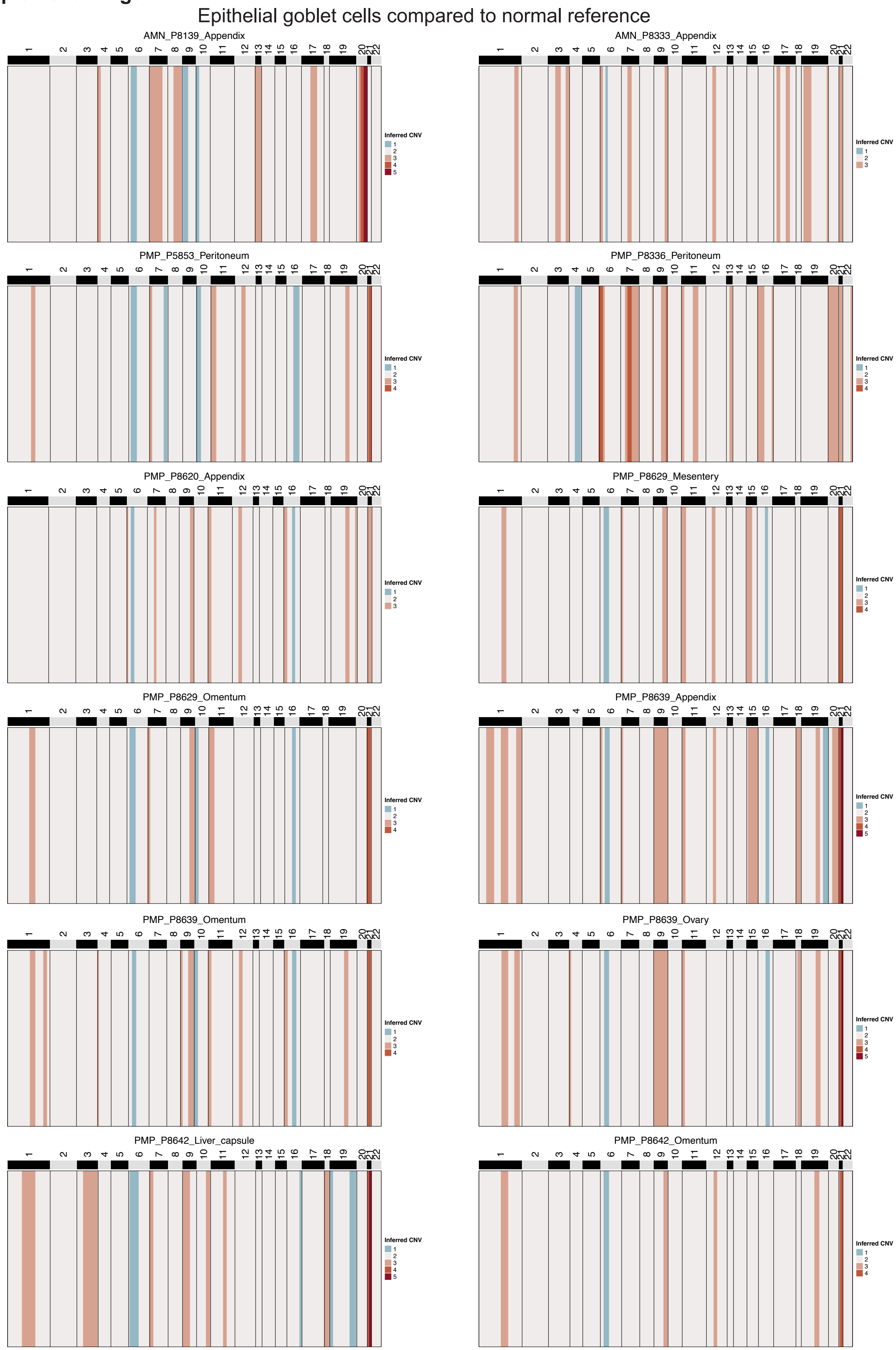

**Supplemental Figure 3**

**A**

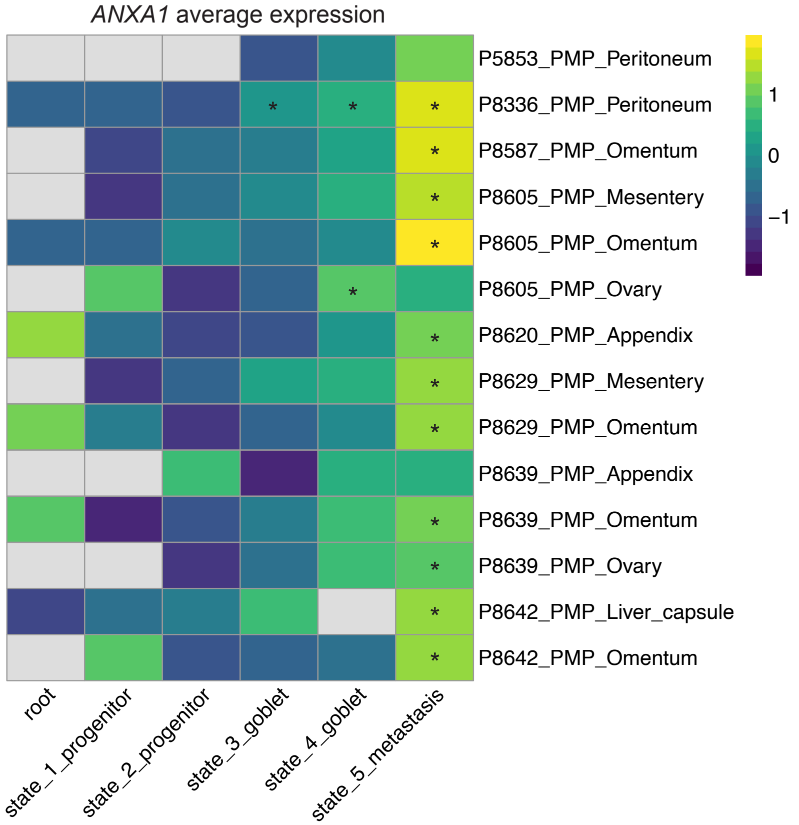


**B**

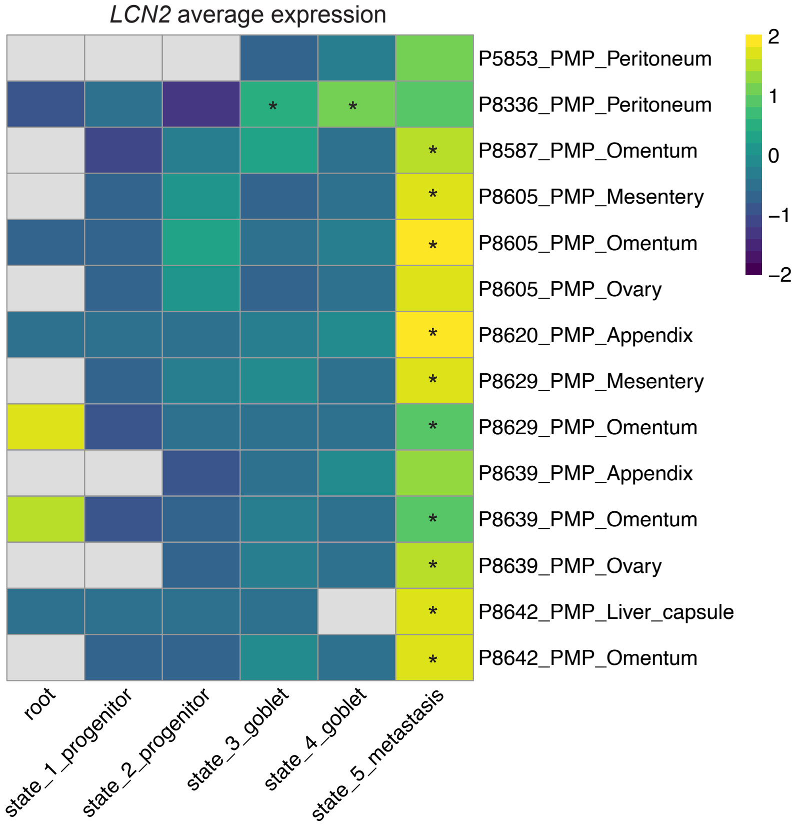

### Supplemental Figure 4

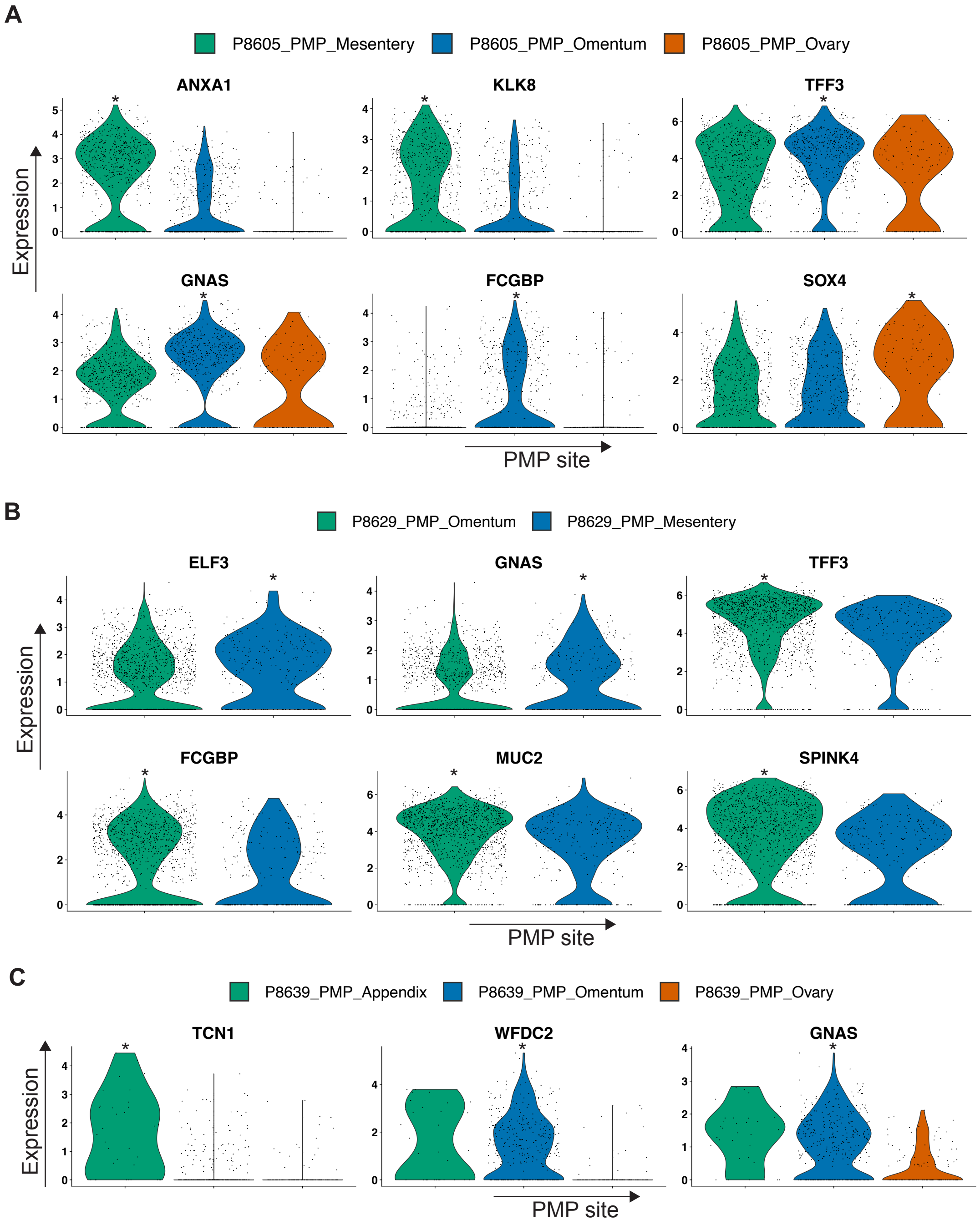
